# Mitochondrial DNA Breaks Activate an Integrated Stress Response to Reestablish Homeostasis

**DOI:** 10.1101/2022.10.01.510431

**Authors:** Yi Fu, Olivia Sacco, Evgeny Kanshin, Beatrix Ueberheide, Agnel Sfeir

**Affiliations:** Skirball Institute of Biomolecular Medicine, Cell Biology Department, NYU School of Medicine, New York, NY, 10016; Molecular Biology Program, Sloan Kettering Institute, Memorial Sloan Kettering Cancer Center, New York, NY, 10065; Proteomics Laboratory, NYU School of Medicine, New York, NY, 10016; Biochemistry and Molecular Pharmacology, NYU School of Medicine, New York, NY, 10016; Department of Neurology, NYU School of Medicine, New York, NY, 10016; Perlmutter Cancer Center, NYU School of Medicine, New York, NY, 10016

## Abstract

Double-strand breaks in mitochondrial DNA (mtDSBs) lead to the degradation of the circular genomes and a reduction in copy number. However, it is unclear how mtDSBs are sensed and what signaling pathways are activated in response to mtDNA damage. In this study, we used mitochondrial-targeted restriction enzymes to investigate the cellular response to mtDSBs. Our results showed that a subset of cells with mtDSBs exhibited defects in mitochondrial protein import, reduced respiratory complexes, and loss of membrane potential. Electron microscopy revealed compromised mitochondrial membrane and cristae ultrastructure. We also found that mtDSBs activated the integrated stress response (ISR) through the phosphorylation of eIF2α by DELE1 and HRI. Notably, inhibition of the ISR exacerbated the mitochondrial import defect and delayed the recovery of mtDNA copy number following break formation. These findings suggest that the ISR plays a role in mitigating mitochondrial dysfunction following mtDNA damage and is critical to promoting mtDNA repopulation. Last, we used proteomics to survey the proteins present in the nucleoids shortly after mtDSBs and identified ATAD3A, a membrane-anchored protein interacting with nucleoids, as a potential factor in transmitting the signal from damaged genomes to the inner mitochondrial membrane. In summary, our study reveals the sequence of events linking damaged mitochondrial genomes with the cytoplasm and highlights the unexpected role of the ISR in reestablishing homeostasis in response to mitochondrial genome instability.

## Introduction

Double-strand breaks in mtDNA (mtDSBs), including ones induced by exogenous and endogenous sources (Alexeyev et al., 2013), lead to linearization and degradation of the circular genomes (Moretton et al., 2017; Srivastava and Moraes, 2001). On the contrary, repair of mtDSBs is rare and results in mtDNA deletions (Phillips et al., 2017), which can cause severe mitochondrial pathologies such as Pearson’s syndrome, Kearns-Sayre syndrome, and chronic progressive external ophthalmoplegia (Tuppen et al., 2010). Degradation of linear mtDNA is primarily carried out by mitochondrial exonuclease activities (MGME1 and POLG), resulting in the loss of mtDNA copy number (Nissanka et al., 2018; Peeva et al., 2018). Replicating residual intact genomes is necessary to replenish mtDNA and requires proper communication between the mitochondria and the nucleus. However, how the cell senses mtDSBs remains poorly understood.

The mitochondria and the nucleus communicate through retrograde signaling, which involves direct inter-organellar contacts, small messenger metabolites, and transcriptional and translational programs (Mottis et al., 2019). Initially discovered in yeast, retrograde signaling involves a group of RTGs that relocate from the mitochondria to the nucleus in response to mitochondrial respiration deficiency and turn on metabolic genes (Butow and Avadhani, 2004). RTG genes are not conserved in higher eukaryotes, and in mammals, one way of communication involves the mitochondrial unfolded protein response (UPRmt) and the integrated stress response (ISR). The ISR is activated through the phosphorylation of a protein called eukaryotic translation initiation factor 2α (eIF2α) by four different kinases in response to various types of stress (Eckl et al., 2021). It reduces global translation but increases the production of specific stress-related proteins, including CHOP, ATF4, and ATF5 (Costa-Mattioli and Walter, 2020). Another form of retrograde signaling involves the release of mitochondrial nucleic acids into the cytoplasm, activating pattern recognition receptors and the innate immune response (Bahat et al., 2021; Dhir et al., 2018; Kim et al., 2019; Yu et al., 2020; Zhong et al., 2018). Activation of an innate immune response also occurs when mtDNA experiences DSBs, resulting in the RIG-I-induced transcription of interferon-stimulated genes without significantly impairing mitochondrial function (Tigano et al., 2021).

Here, we uncover the cellular response to mtDNA damage. We observed severe dysfunction in a subset of cells with mtDSBs, including loss of membrane potential, defective protein import, and abnormal cristae morphology. Furthermore, cells with mtDNA breaks activate the ISR in a DELE1-HRI-dependent manner (Fessler et al., 2020; Guo et al., 2020). We show that activating an acute ISR is crucial for mtDNA repopulation after damage and restoring mitochondrial function. We also identify ATAD3A, an AAA+ ATPase located at the inner mitochondrial membrane, as a factor that potentially signals from defective genomes to mitochondrial membranes. Our findings provide insight into how cells mitigate mitochondrial dysfunction in response to mtDNA instability.

## Results

### Mistargeting of mitochondrial-targeted proteins in a subset of cells with mtDNA breaks

To trigger DSBs in mtDNA, we used the previously established mitochondrial-targeted restriction enzyme ApaLI that cleaves in the MT-RNR1 locus encoding 12S rRNA (Bayona-Bafaluy et al., 2005). We expressed doxycycline (Dox) inducible mito-ApaLI and confirmed the accumulation of cleaved mtDNA 12 hours after Dox treatment (Figures 1A and 1B). As a control, we expressed an inactive mito-ApaLI (mito-ApaLI*) comprising three base substitutions at the predicted catalytically active site (D211DA, E225A, K227A). When expressed in 293T cells, mito-ApaLI* did not cleave mtDNA or cause depletion of mtDNA copy number (Orlowski and Bujnicki, 2008) (Figure S1A and S1B).

**Figure 1.**
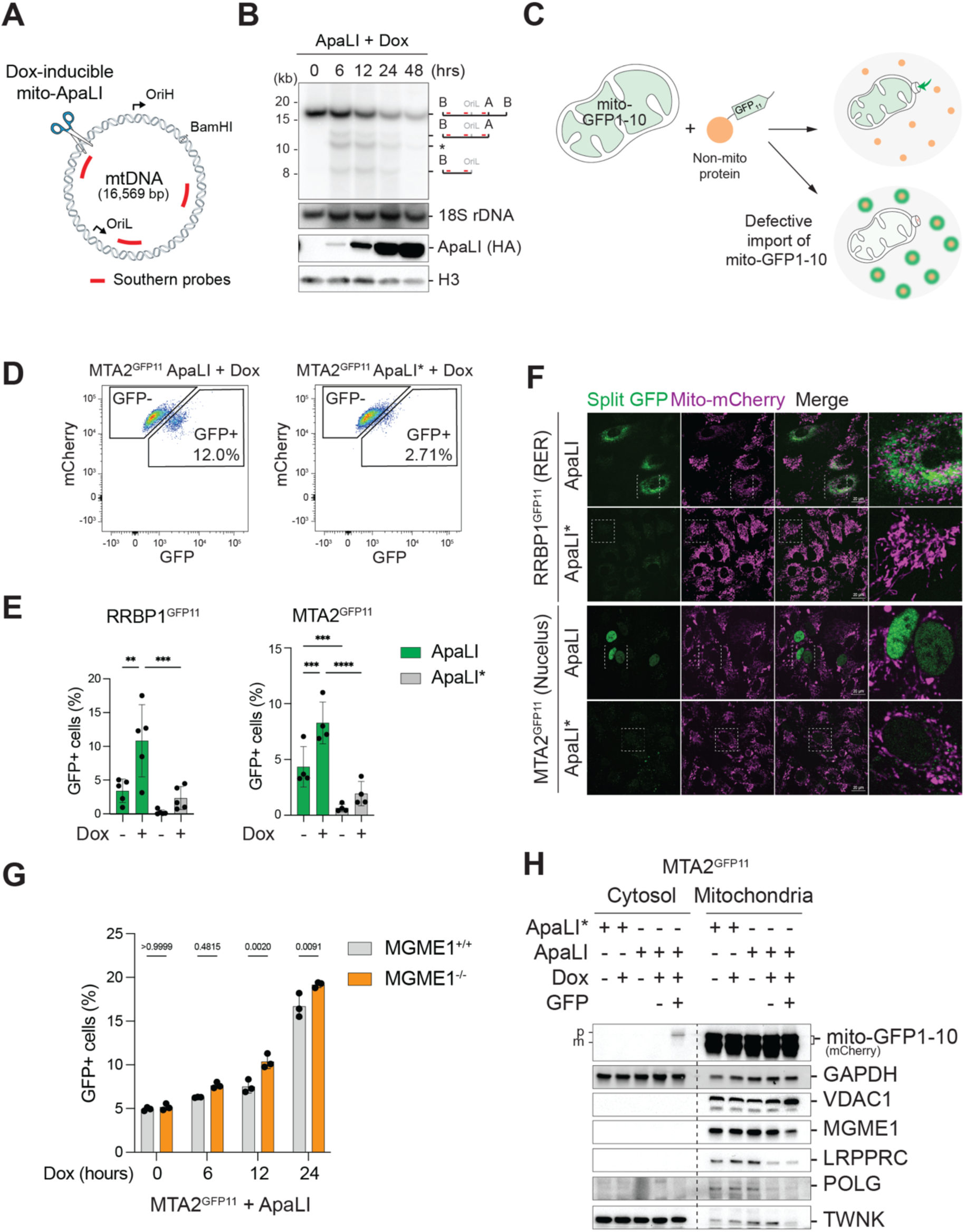
Impaired mitochondrial protein import in response to mtDNA breaks. (A) Schematic of introducing mtDSB using a mitochondria-localized restriction enzyme mito-ApaLI controlled by a doxycycline (Dox) inducible promoter. Depicted in red are probes used in the southern blot recognizing different mtDNA regions. Extracted mtDNA was linearized with BamHI retriction enzyme. OriH, the origin of heavy-strand replication; OriL, the origin of light-strand replication. (B) Southern blot analysis of mtDNA and nuclear 18S rDNA (loading control) and western blot analysis of mito-ApaLI (anti-HA) and histone H3 (loading control) in cells expressing Dox-induced mito-ApaLI for 0-48 hours. A, ApaLI restriction site; B, BamHI restriction site; (*) depicts a degradation intermediate. (C) Schematic of the split-GFP system used to assess mitochondrial protein import following mtDSBs. A mitochondrial localization signal targets mito-GFP1-10 to the mitochondrial matrix. Non-mitochondrial proteins were tagged endogenously with four tandem GFP11 repeats using CRISPR/Cas9 editing. Defective mitochondrial protein import leading to mislocalization of mito-GFP1-10 would lead to GFP reconstitution outside the mitochondria where GFP11 resides. (D) Representative gating strategy for fluorescence-activated cell sorting (FACS) of GFP negative (GFP-) and positive (GFP+) populations from MTA2^GFP11^ cells expressing Dox-induced mito-ApaLI and catalytically inactive mito-ApaLI* for two days. (E) Percentage of GFP+ cells expressing mito-GFP1-10 and GFP11-tagged RRBP1 (RRBP1^GFP11^) and MTA2 (MTA2^GFP11^) subjected to mtDSBs. RRBP1^GFP11^ and MTA2^GFP11^ cells expressing Dox-inducible mito-ApaLI and mito-ApaLI* were treated with Dox for two days. (F) Representative images depicting the subcellular localization of reconstituted split-GFP in RRBP1^GFP11^ and MTA2^GFP11^ cells used in (E). RER, rough endoplasmic reticulum. Scale bar, 20 μm. (G) Graph representing the GFP+ percentage in wildtype and MGME1-depleted MTA2^GFP11^ cells expressing Dox-induced mito-ApaLI for 0, 6, 12, and 24 hours, analyzed by flow cytometry. (H) Subcellular fractionation followed by western blot analysis of the indicated proteins in MTA2^GFP11^ cells with the indicated treatments. GFP, FACS isolated GFP- or GFP+ cells. p, precursor isoform; m, mature isoform. See also Figure S3A. See also Figure S1, S2, and S3.

Mitochondrial protein import, which supplies the majority of mitochondrial proteome and ensures balanced composition of the electron transport chain, serves as a signal for mitochondrial dysfunction (Song et al., 2021). We used a split-GFP system (Leonetti et al., 2016) to investigate potential defects in mitochondrial protein import in response to mtDSBs. Specifically, a truncated form of GFP (GFP1-10) was targeted to the mitochondria using a mitochondrial localization sequence (MLS). A separate fragment encoding the GFP11 peptide was attached to abundant non-mitochondrial proteins (Figure 1C). Mistargeted mito-GFP1-10 will be sequestration in the cytoplasm and associated with GFP11. This would then reconstitutes the GFP signal in the cytoplasm, which can be detected using flow cytometry and fluorescent microscopy. As a control, we appended GFP11 to the C-terminus of TFAM and LRPPRC and confirmed the proper localization of mito-GFP1-10 to the mitochondrial matrix in the absence of mitochondrial DNA breaks (Figure S1C). To probe for mistargeting of mito-GFP1-10 outside of the mitochondria, we targeted GFP11 using CRISPR/Cas9 to the C-terminus of proteins located in the rough endoplasmic reticulum (RRBP1) (Shibata et al., 2010) and the nucleus (MTA2 and SMCHD1) and generated clonal cells with successful knock-in (termed RRBP1^GFP11^, MTA2^GFP11^, and SMCHD1^GFP11^ cells) (Figure S1D). Overexpression of mito-ApaLI* in MTA2^GFP11^ cells resulted in ∼2% of GFP-positive (GFP+) cells (Figure 1E). This result can be explained by the obstruction of the mitochondrial import machinery previously observed upon overexpression of mitochondrial proteins (Weidberg and Amon, 2018). While leaky expression of mito-ApaLI led to a small but significant fraction of GFP-positive cells, Dox-induced mito-ApaLI expression led to a significant increase in the GFP+ percentage (up to 20% GFP+ cells within 48 hours of Dox), highlighting substantial import defects due to mtDSBs (Figure 1D-E). Using fluorescent microscopy, we confirmed that the GFP signal was in the rough endoplasmic reticulum in RRBP1^GFP11^ cells and the nucleus in MTA^GFP11^ and SMCHD1^GFP11^ cells (Figure 1F, S1E and S1F). We corroborated these findings by targeting GFP11 to additional non-mitochondrial proteins, including ER (KTN1, VAPB, CANX, SEC61B) and Golgi (ARL1) residing proteins, and observed similar accumulation of GFP signal outside of the mitochondria upon mito-ApaLI-driven mtDSBs (Figure S1D and S1G). Based on these results, we concluded that in a subset of cells with mtDNA breaks, mislocalization of mitochondrial GFP1-10 promotes its assembly with GFP11-tagged proteins in the cytoplasm and the nucleus.

Given that cleaved mtDNA molecules are ultimately degraded (Moretton *et al*., 2017; Srivastava and Moraes, 2001), we speculated that the mislocalization of mitochondrial-targeted proteins could result from the loss of mtDNA copy number. To test this possibility, we depleted mtDNA by treating MTA^GFP11^ cells with chain terminator 2′,3′-dideoxycytidine (ddC) for up to 6 days, which led to a reduction in mtDNA copy number that was equivalent to 48 hours of ApaLI expression (Figure S2A). We noted that the accumulation of GFP+ cells was significantly more in response to mtDNA breaks than following the depletion of mtDNA (Figure S2B). These results are consistent with mtDNA breaks specifically triggering a mitochondrial import defect. To corroborate these findings, we blocked mtDNA degradation following break formation by deleting the exonuclease MGME1 (Figure 1G, S2C, and S2D). Quantification of copy number by qPCR confirmed the persistence of cleaved mtDNA in MGME1^-/-^ cells 24 hours after mito-ApaLI induction (Figure S2D). Despite the persistence of linear mtDNA molecules and mitochondrial encoded proteins (i.e., COX1 and ND2) in the MGME1^-/-^ cells expressing mito-ApaLI (Figure S2F), we observed significant mitochondrial protein mislocalization measured by cytoplasmic trapping of mito-GFP1-10 (Figure 1G). Taken together, these observations suggest that mitochondrial defects in response to mito-ApaLI are triggered by broken mtDNA molecules and not driven by subsequent depletion of mtDNA.

The mistargeting of mito-GFP1-10 is consistent with defects in mitochondrial protein import in response to mtDSBs. To examine the subcellular localization of mitochondrial-targeted proteins, we sorted cells based on GFP positivity and performed fractionation experiments followed by western blot analysis. We detected mito-GFP1-10 precursor in the cytosolic fraction in the GFP+ cells but not in the GFP-cells and other control samples (Figure 1H and S3A). Furthermore, when we overexpressed frataxin, which has distinct isoforms indicative of mitochondrial import and processing (Branda et al., 1999; Nabhan et al., 2015), cytosolic precursors were enriched in the presence of mtDSBs (Figure S3B and S3C). Last, we observed a significant reduction in the endogenous levels of several nuclear-encoded mitochondrial proteins in the GFP+ cells following mtDSBs, including POLG, Twinkle, MGME1, and LRPPRC (Figure 1H and S3A). Altogether, our data suggest a deficiency in mitochondrial protein import in response to mtDSBs.

### mtDNA damage leads to defective mitochondrial membrane ultrastructure

We suspected that mistargeting of mitochondrial proteins represents broader mitochondrial dysfunction in a subset of cells experiencing mtDNA breaks. To test this, we induced mtDNA breaks in RRBP1^GFP11^ cells, isolated the GFP+ population, and measured various indices of mitochondrial function, including membrane potential, levels of proteins involved in oxidative phosphorylation (OXPHOS), and mitochondrial morphology. Staining with tetramethylrhodamine methyl ester (TMRM) revealed a significant loss of membrane potential in GFP+ cells relative to GFP-cells and unsorted cells expressing mito-ApaLI or mito-ApaLI* (Figure 2A, 2B, S4A and S4B). Furthermore, GFP+ cells expressing mito-ApaLI* did not show loss of membrane potential, further suggesting that mitochondrial defects inherent to mtDSBs are not common to other mitochondrial stressors, such as protein overload. We observed a decrease in OXPHOS components upon mtDNA break induction, which was more pronounced in GFP+ cells (Figure S4C). Additionally, GFP+ cells with mtDSBs displayed increased autophagy, as determined by monitoring the conversion of LC3-I to LC3-II (Figure S4D). In conclusion, our results highlight significant mitochondrial dysfunction in a subset of cells with mtDNA breaks.

**Figure 2.**
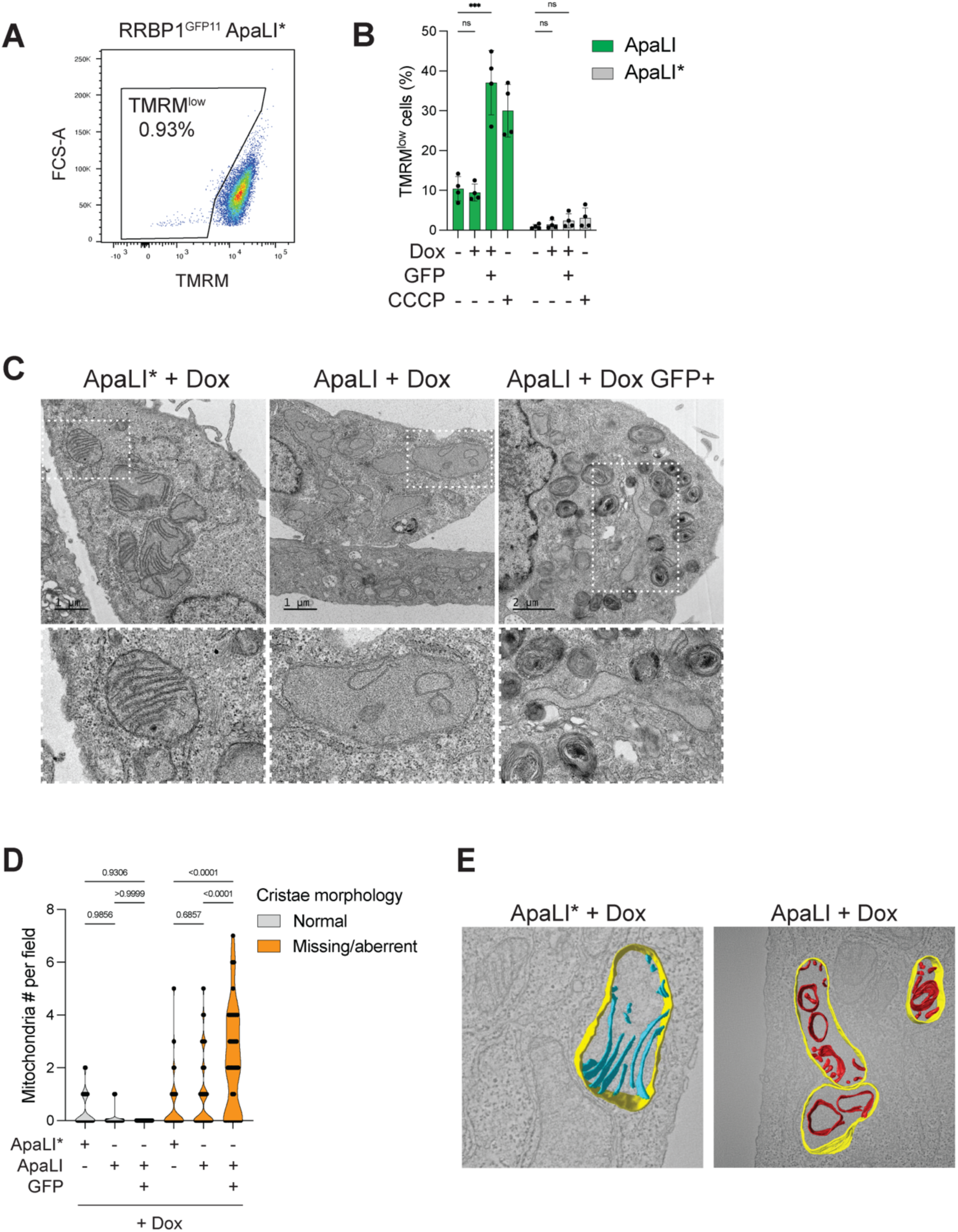
Defective mitochondrial membranes in cells subjected to mtDNA damage. (A) Gating strategy to quantify the percentage of cells with loss of membrane potential based on the loss of TMRM staining signal (TMRM^low^ cells). Representative flow cytometry analysis in uninduced mito-ApaLI* RRBP1^GFP11^ cells. (B) Flow cytometry analyzed the percentage of TMRM^low^ population in RRBP1^GFP11^ cells in response to mito-ApaLI mtDSBs. Mitochondrial uncoupler CCCP (Carbonyl cyanide m-chlorophenyl hydrazone) was used at 20 μM for 6 hours. Gating was based on uninduced mito-ApaLI* cells as in (A). Data are means ± s.d. of *n* = 4 biological replicates; two-way ANOVA. \**P* ≤ 0.05, \*\**P* ≤ 0.01, \*\*\**P* ≤ 0.001, \*\*\*\**P* ≤ 0.0001. (C) Representative transmission electron microscopy (TEM) images of unsorted and GFP+ MTA2^GFP11^ cells expressing Dox-induced mito-ApaLI (and mito-ApaLI*) for three days. (D) Violin plot depicting numbers of normal and abnormal mitochondria per TEM field. Total fields analyzed for mito-ApaLI*+Dox, mito-ApaLI+Dox, and mito-ApaLI+Dox GFP+ samples are 35, 48, and 34, respectively. Two-way ANOVA. (E) Representative 3D reconstructed mitochondria from TEM images of MTA2^GFP11^ cells expressing Dox-induced mito-ApaLI* and mito-ApaLI for three days. See also Figure S4 and Video 1-5.

We next used transmission electron microscopy (TEM) to examine the mitochondria structure in cells expressing mito-ApaLI and noticed significant structural deformation. Specifically, the GFP+ subpopulation showed severe membrane abnormalities and loss of cristae structures (Figure 2C and 2D; Video 1-3). In addition, GFP+ cells accumulated multi-membrane lysosomes (Wong et al., 2018), consistent with increased LC3 conversion (Figure S4D and S4E). 3D reconstruction of the TEM images revealed that the mitochondria in cells under mtDSBs lacked the typical lamellar structures and had no cristae junctions but vesicular and onion-shaped cristae (Figure 2E; Video 4 and 5), which are similar to structures observed when the MICOS complex is depleted (Stephan et al., 2020). We also saw a significant loss of MIC60 and MIC10, two critical components of the MICOS complex, in the GFP+ cells with mtDNA breaks (Figure S4F). Overall, significant abnormalities in the structure of the mitochondria correlate with severe mitochondrial dysfunction in response to mtDNA breaks.

### DELE1–HRI activates a robust integrated stress response in cells with mtDSBs

So far, our data revealed severe defects in mitochondria in a subset of cells experiencing mtDNA breaks. To examine the broader cellular response to mtDNA damage, we performed RNA-seq analysis. We compared the transcriptomes of sorted GFP+ RRBP1^GFP11^ and MTA2^GFP11^ cells to control cells, including the GFP-subpopulation and unsorted cells expressing mito-ApaLI and mito-ApaLI* (Figure 3A-D and Figure S5A-B). We found that 558 genes were deregulated in GFP+ cells compared to GFP-cells, with the latter showing a similar transcriptional profile to the unsorted cells (Figure 3A). Gene Set Enrichment Analysis (GSEA) revealed several biological processes that were significantly altered, including upregulation of the inflammatory response pathway and downregulation of mitotic genes (Figure 3B and S5A). We also observed induction of the unfolded protein response (UPR) pathway in GFP+ cells compared to control cells (Figure 3B).

**Figure 3.**
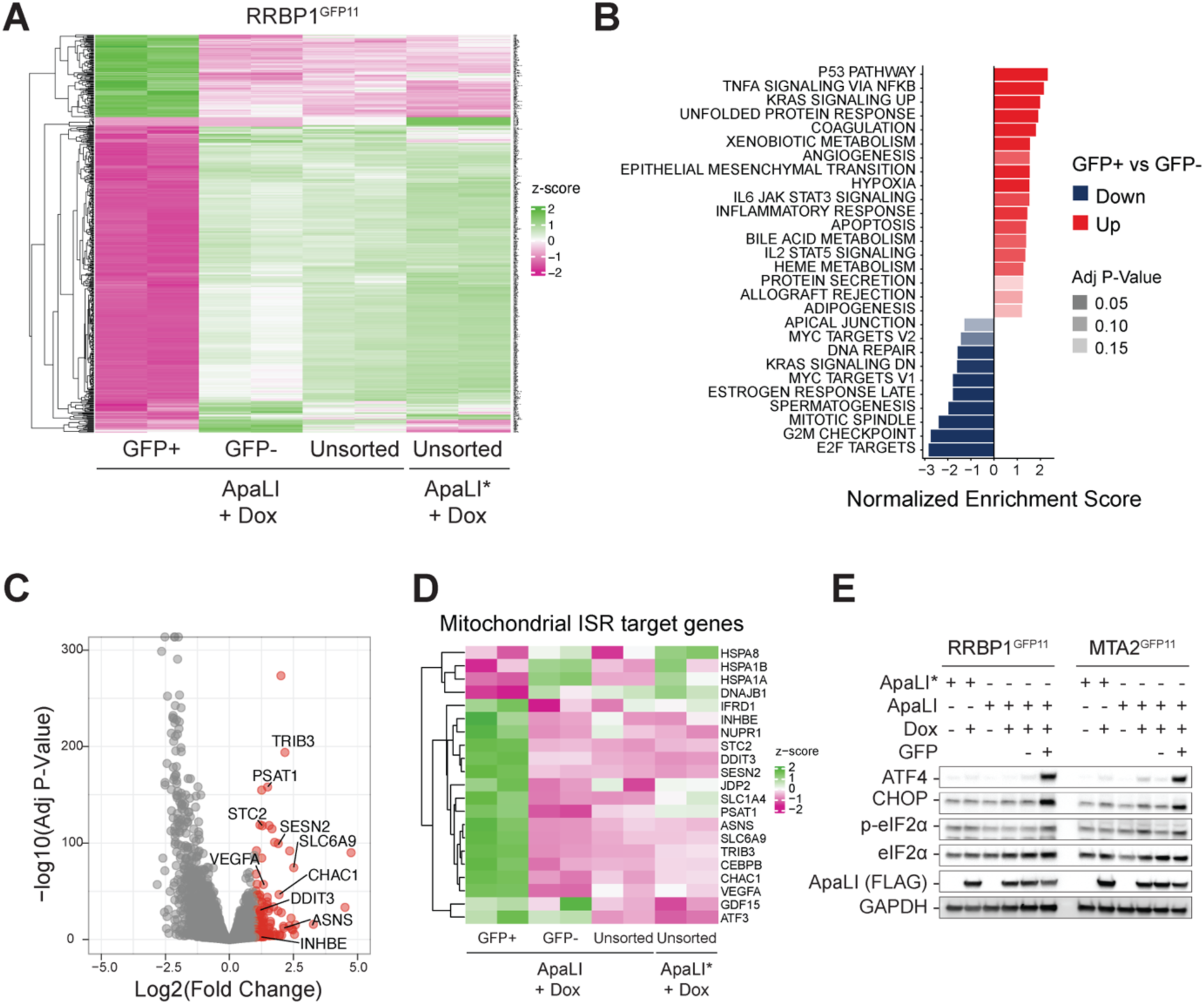
Activation of the integrated stress response in a subset of cells under mtDSBs. (A) Heatmap depicting expression of 558 differentially expressed genes in GFP+ versus GFP-RRBP1^GFP11^ cells with mito-ApaLI induced for three days (116 up-regulated, 442 down-regulated. 2 fold change cut-off, adjusted p-value < 0.01, *n* = 2 biological replicates). (B) Gene Set Enrichment Analysis of differentially expressed genes depicted in (A) using MSigDB hallmark gene sets. Red bars indicate up-regulated pathways, and blue bars represent down-regulated genes in GFP+ cells compared to GFP-cells. (C) Volcano plot highlighting 116 up-regulated genes in GFP+ relative to GFP-subpopulations in red. ISR target genes are labeled with gene symbols. (D) Heatmap showing the expression of mitochondrial ISR target genes. The list of ISR target genes specific to mitochondrial dysfunction was published previously (Fessler et al., 2020). (E) Western blot analysis of ATF4, CHOP, and phosphorylated eIF2α in GFP+ population of mito-ApaLI induced RRBP1^GFP11^ and MTA2^GFP11^ cells, compared to the GFP- and unsorted cells. See also Figure S5.

The integrated stress response (ISR) plays a central role in the mitochondrial UPR in mammalian cells (Anderson and Haynes, 2020). Consistent with this, we observed increased expression of many ISR target genes (Figure 3C, 3D, and S5B) and accumulation of ATF4 and CHOP (Figure 3E) in GFP+ cells in response to mito-ApaLI induction. The accumulation of ATF4 and CHOP was specific to GFP+ cells in response to mtDNA breaks and was significantly higher than levels observed in GFP+ population of cells expressing the inactive mito-ApaLI* (Figure S5C and S5D). Furthermore, mtDNA depletion with ddC treatment did not activate a robust ISR response (Figure S5E and S5F). These data further support that the activation of the ISR in response to mtDNA breaks is inherent to the presence of cleaved DNA molecules.

The phosphorylation of eIF2α has been linked to four kinases: HRI, PKR, PERK, and GCN2, each activated by different stress stimuli (Anderson and Haynes, 2020; Costa-Mattioli and Walter, 2020). Previous research has shown that inhibition of OXPHOS and membrane depolarization can activate the inner membrane protease OMA1, which cleaves and triggers relocation of DELE1 from the intermembrane space to the cytosol, where it activates HRI (Fessler et al., 2020; Guo et al., 2020). To identify the kinase that phosphorylates eIF2α in response to mtDNA breaks, we individually depleted HRI, PKR, PERK, and GCN2 in MTA2^GFP11^ cells using CRISPR/Cas9, induced the expression of mito-ApaLI, and then measured levels of CHOP and ATF4 in the sorted GFP+ cells. Depletion of HRI, but not PKR, PERK, or GCN2, led to a reduction in the activation of the ISR (Figure 4A, S6A-B). Furthermore, inhibition of DELE1, the upstream activator of HRI, had a similar effect (Figure 4A, S6A-B). In contrast, our data indicate that OMA1 – previously shown to respond to ATP synthase inhibition, mitochondrial depolarization, and cardiomyopathy (Ahola et al., 2022; Fessler *et al*., 2020; Guo *et al*., 2020) – is dispensable for eIF2α phosphorylation in response to mtDNA damage (Figure 4A, S6A, and S6B). In conclusion, the ISR is activated in a DELE1-HRI-dependent manner in a subset of cells experiencing mtDNA breaks.

**Figure 4.**
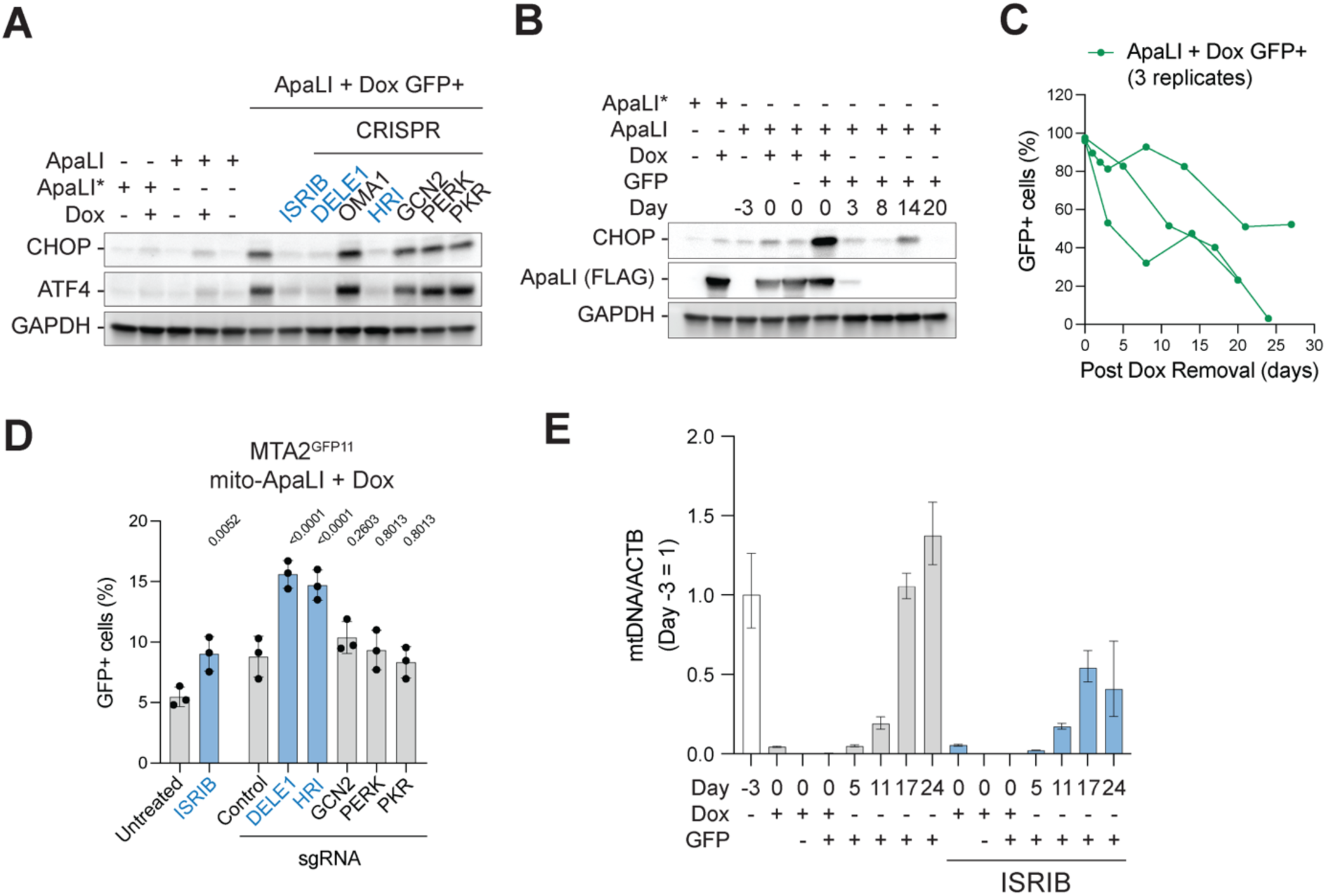
DELE1-HRI triggers a robust ISR upon mtDSBs to restore homeostasis. (A) Western blot analysis of CHOP and ATF4 in the GFP+ population of MTA2^GFP11^ cells with mtDSBs and subjected to ISR inhibition using ISRIB, and CRISPR depletion of DELE1, OMA1, HRI, GCN2, PERK, and PKR. See Figure S6A for knock-out efficiencies and Figure S6B for western blot quantification. (B) Temporal recovery of CHOP levels in sorted GFP+ population of MTA2^GFP11^ cells in the absence of Dox, analyzed by western blots. (C) Temporal dynamics of GFP+ percentage after sorting and Dox removal, analyzed by flow cytometry. Figure S6C represents GFP dynamics in unsorted cells. (D) Percentage of GFP+ MTA2^GFP11^ cells under mtDSBs and subjected to indicated ISR inhibition treatments, analyzed by flow cytometry. Data are means ± s.d. of n = 3 biological replicates: one-way ANOVA. The P-value for ISRIB treatment was calculated by comparing it to the Untreated. P-values for CRISPR treatments were calculated by comparing them to the Control (non-targeted sgRNA). See Figure S6D for CRISPR editing efficiencies. (E) Copy number dynamics of mtDNA in sorted GFP+ populations of MTA2^GFP11^ cells with the indicated treatments. MTA2^GFP11^ cells expressing mito-ApaLI were treated with Dox on Day -3, and Dox was withdrawn on Day 0. Copy numbers were analyzed by qPCR using primers aginst MT-RNR1/2 (mtDNA) relative to ACTB. Values at different time points were normalized to Day - 3. Normalized copy numbers mean ± s.d. from *n* = 3 technical replicates. See Figure S6G for additional biological replicates. See also Figure S6.

### Activation of the ISR counteracts mitochondrial dysfunction and promotes mtDNA repopulation in response to mtDSBs

Acute activation of the ISR is typically an adaptive response that allows cells to recover from stress, while prolonged ISR can lead to cell death (Koromilas, 2019). To determine whether the activated ISR in response to mtDNA damage is reversible, we withdrew Dox from sorted mito-ApaLI-induced GFP+ cells and monitored the accumulation of CHOP over time. Our data indicated that CHOP levels dropped within three days of Dox withdrawal (Figure 4B), suggesting an acute and reversible ISR response following mtDNA breaks. We observed a concomitant decrease in the percentage of GFP+ cells following Dox withdrawal (Figure 4C and S6C).

To understand the impact of the ISR on cells experiencing mtDNA breaks, we treated MTA2^GFP11^ cells expressing mito-AapLI with ISRIB, a small molecule inhibitor of the ISR (Costa-Mattioli and Walter, 2020; Sidrauski et al., 2015; Tsai et al., 2018; Zyryanova et al., 2018). Using the reconstitution of GFP as a proxy for mitochondrial protein mislocalization in MTA2^GFP11^ Cells, we found that treatment with ISRIB led to a significant increase in the percentage of GFP+ cells in response to mtDNA breaks (Figure 4D). We obtained similar results when we depleted HRI and DELE1 using CRISPR-Cas9, suggesting that the ISR counteracts defective mitochondrial protein import in response to mtDNA breaks (Figure 4D and S6D). Since the mitochondria lack DSB repair activity, mtDNA DSBs, including those induced by mito-ApaLl, result in rapid degradation of the genomes and ultimately lead to a loss of mtDNA copy number (Figure S6E and S6F). However, after the restriction enzyme is turned off, mtDNA genomes are repopulated to restore cellular homeostasis (Figure S6E and S6F). Interstingly, inhibition of the ISR delayed the recovery of mtDNA copy number relative to non-treated cells (Figure 4D and S6G). In conclusion, we propose that ISR exerts a pro-survival and adaptive role in counteracting the defects caused by mtDNA damage, enabling mtDNA copy number recovery and restoration of cellular homeostasis.

### BioID-based proteomic profiling reveals changes in nucleoid composition in response to mtDSBs

Having uncovered the downstream impact of mtDSBs, we next turned our attention to upstream factors within the mitochondrial matrix and the inner mitochondrial membrane that could sense mtDNA damage. To identify proteins that interact with mtDNA in the presence of DSBs, we used a proteomics-based approach that combines proximity-dependent biotin labeling with quantitative proteomics. We transduced cells with a fusion protein of TFAM and BioID2 (Kim et al., 2016), which localized to the mitochondria without affecting mtDNA copy number (Figure 5A, S7A, and S7B). We then cultured the cells with heavy, medium, and light isotopes of lysine and arginine for ten population doublings and treated cells with doxycycline to induce the expression of mito-ApaLI or mito-ApaLI*, and biotin to trigger the biotinylation of proteins adjacent to TFAM-BioID2 (Figure 5B and S7C). Samples were collected 12 hours after treatment, and the biotinylated proteins were captured with streptavidin beads and analyzed using mass spectrometry (Figure 5B). This approach allowed us to identify proteins that interact with mtDNA in the presence of DSBs and potentially play a role in the cellular response to mtDNA damage.

**Figure 5.**
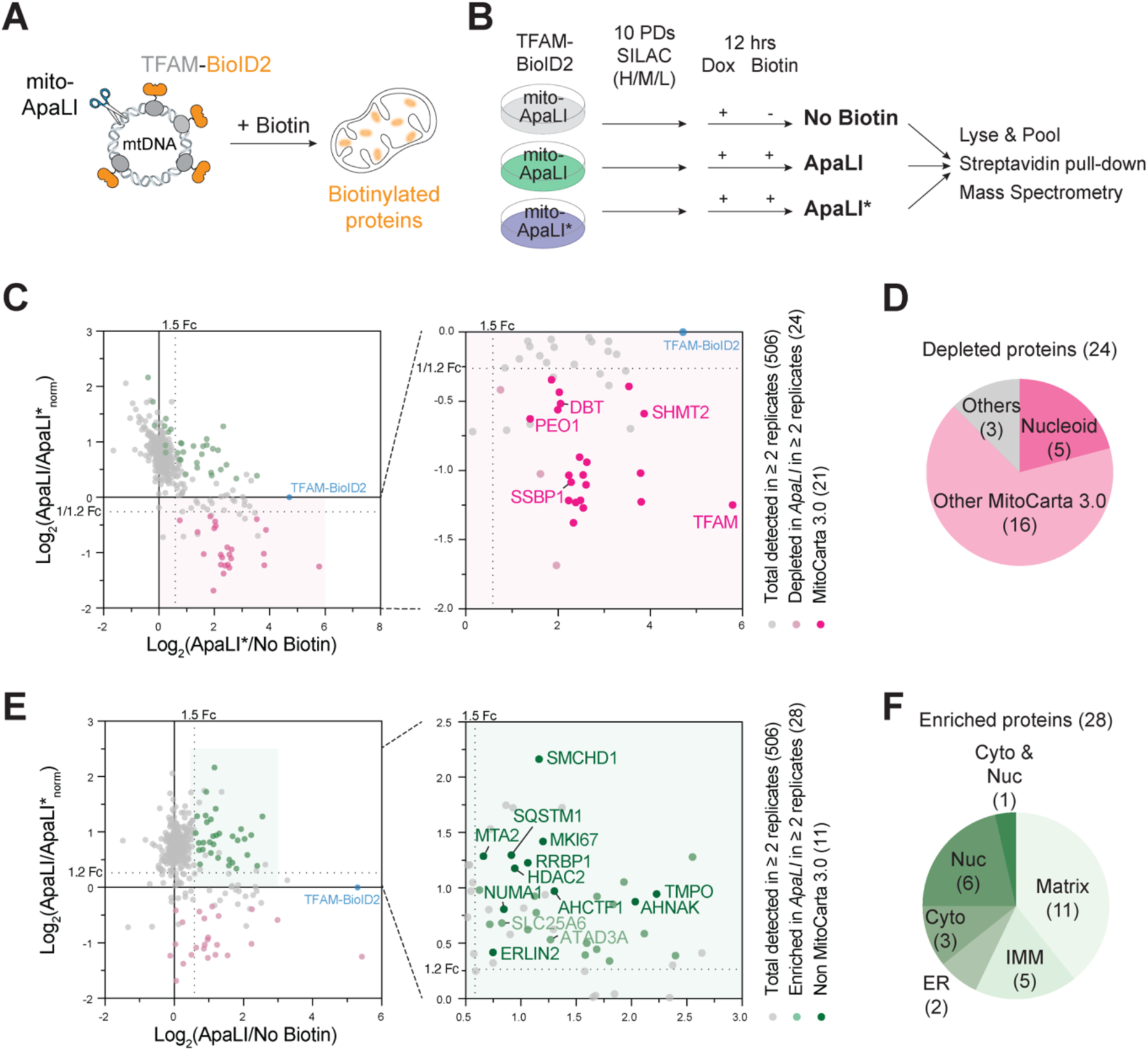
BioID-based proteomic analysis reveals changes in nucleoid composition in response to mtDSBs. (A) Scheme of TFAM-BioID2 biotinylation to capture mtDNA-associated proteins. (B) Experimental design of a 3-state SILAC labeling used to identify mtDNA-associated proteins in response to mtDSBs. H/M/L, heavy/medium/light isotopes of lysine and arginine. PD, population doubling. (C) SILAC ratio plot showing the TFAM-BioID2-normalized log2(ApaLI/ApaLI*) against log2(ApaLI*/No Biotin). Twenty-four proteins depleted in mito-ApaLI are highlighted in pink. Among them, 21 proteins belonging to MitoCarta 3.0 are marked in dark pink on the right. See also Figure S2D. (D) Categories of 24 proteins depleted in mito-ApaLI. (E) SILAC ratio plot showing the normalized log2(ApaLI/ApaLI*) against log2(ApaLI/No Biotin). Twenty-eight proteins enriched in ApaLI are highlighted in green. Among them, 11 proteins that do not belong to MitoCarta 3.0 are marked in dark green on the right. (F) Categories of subcellular localization of 28 mtDSB-enriched proteins. Matrix, mitochondrial matrix; IMM, inner mitochondrial membrane; ER, endoplasmic reticulum; Cyto, cytoplasm; Nuc, nucleus. See also Figure S7, S8, and Table 1.

We calculated the SILAC ratio (Cox and Mann, 2008) of peptides retrieved from samples treated with biotin (mito-ApaLI and mito-ApaLI*) compared to the control cells not treated with biotin. Our analysis identified 68 and 94 proteins (1.5-fold change, ≥ 2 replicates) that were in the vicinity of nucleoids in cells expressing mito-ApaLI and mito-ApaLI*, respectively (Figure S7D). Most nucleoid proximal factors (>70%) were previously detected in the mitochondria based on MitoCarta 3.0 (Rath et al., 2021) (Figure S8A). Furthermore, several hits were nucleoid-associated proteins as curated in (Han et al., 2017) (Figure S8B). To monitor changes in nucleoid composition following mtDNA breaks, we calculated the SILAC ratio of proteins retrieved in mito-ApaLI relative to mito-ApaLI*\** (Figure S7D and S8C). As expected, nucleoid proteins and several adjacent ribosomal proteins were depleted in response to mtDNA breaks (Figure 5C, 5D, and S8D) (Gilkerson et al., 2013; Xavier and Martinou, 2021). On the other hand, we detected 28 proteins to be enriched in the vicinity of TFAM-BioID2 shortly after break formation. These included cytoplasmic and nuclear proteins, which likely resulted from the mistargeting of the bait to the cytosol upon mtDNA damage (Figure 5E and 5F). Notably, we also identified several mitochondrial matrix proteins and factors associated with the inner mitochondrial membrane (IMM) (Figure 5E, 5F, and S8E) that could potentially act as sensors of mtDNA damage.

### ATAD3A as a potential link between mtDNA damage and defective mitochondrial membranes

Last, we sought to highlight the putative factor that could transmit the signal from defective mitochondrial genomes to the mitochondrial double membranes. To do so, we leveraged our proteomic analysis focusing on proteins enriched in the vicinity of nucleoids carrying mtDNA breaks. We depleted candidate factors, including ATAD3A, DNAJA3, HSPA9, SLC25A6, LRPPRC, and TUFM in MTA2^GFP11^ (and RRBP1^GFP11^) cells. We then induced mito-ApaLI expression and assessed mistargeting of mito-GFP1-10 using flow cytometry. Among the candidates we tested, knockdown of ATAD3A resulted in an increase in GFP+ cells in response to break formation (Figure 6A). ATAD3A is a conserved AAA+ ATPase that regulates mitochondrial inner membrane architecture. It spans the IMM with its C-terminus facing the matrix and associating with the nucleoids (Gilquin et al., 2010; Ishihara et al., 2022; Sen et al., 2022) (Figure S9A). We surmised that ATAD3A might link nucleoids comprising defective genomes with mitochondrial membranes. To corroborate our findings, we performed CRISPR/Cas9 editing to delete ATAD3A in MTA2^GFP11^ cels, and observed synergistic defects in mitochondrial protein mislocalization and ISR activation in the presence of mtDNA breaks (Figure 6B and 6C). In conclusion, our results are consistent with ATAD3A potentially acting downstream of mtDNA damage to alter mitochondrial membrane structure and trigger an ISR.

**Figure 6:**
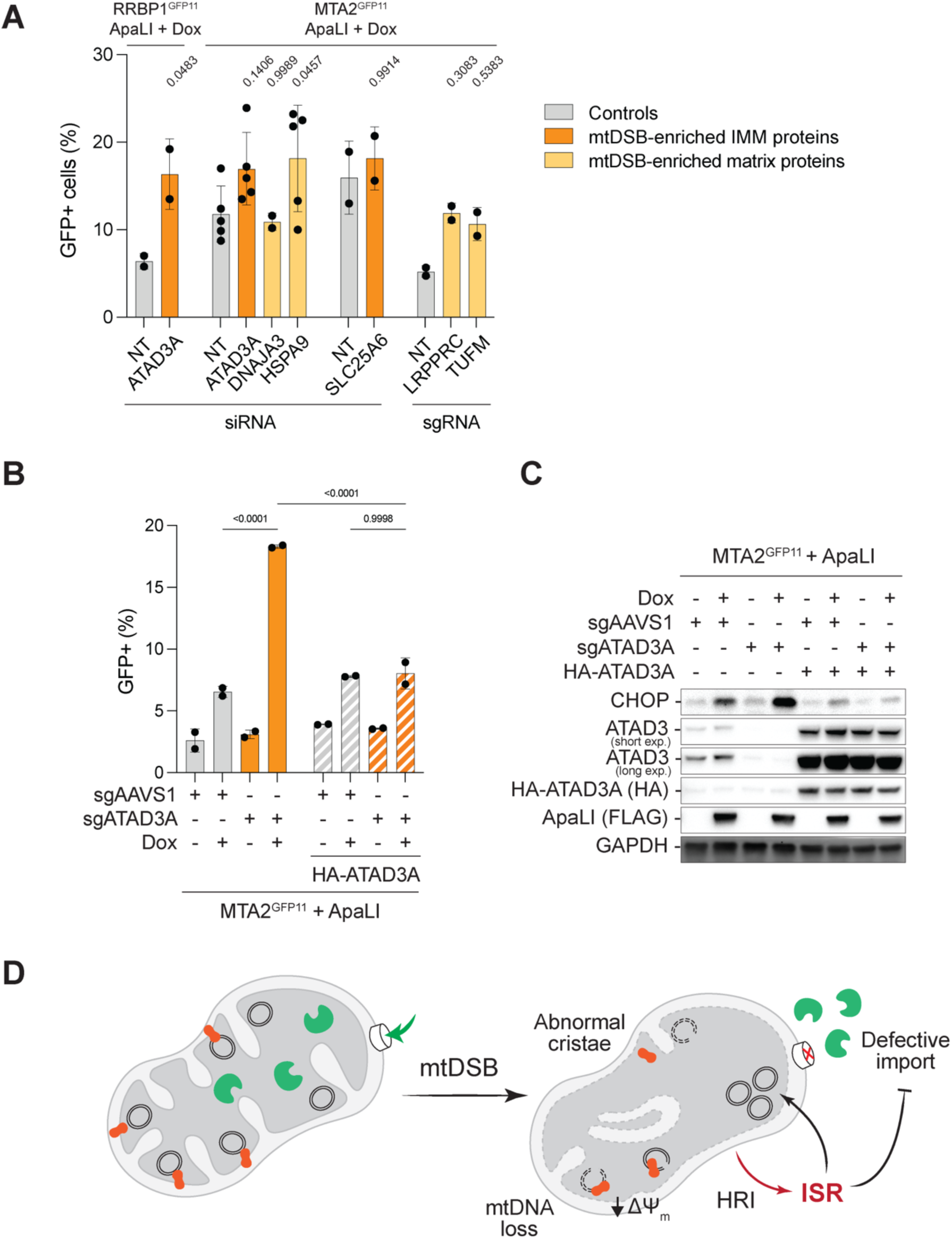
ATAD3A, a potential link between mtDNA damage and the ISR. (A) Percentage of GFP+ cells in MTA2^GFP11^ or RRBP1^GFP11^ cells subjected to siRNA or CRISPR perturbation of candidate genes from the proteomics as indicated, and expressing Dox-induced mito-ApaLI for two days, determined by flow cytometry analysis. Data are means ± s.d. of *n* = 2-4 biological replicates; one-way ANOVA. P-values were calculated by comparing to the corresponding non-targeted (NT) siRNA or sgRNA controls. (B) Percentage of GFP positive cells in MTA2^GFP11^ cells overexpressed with ATAD3A, treated with sgRNA targeting AAVS1 or ATAD3A locus and subjected to Dox-induced mito-ApaLI mtDSBs, determined by flow cytometry analysis. Data are means ± s.d. of *n* = 2 biological replicates; two-way ANOVA. (C) Expression of CHOP and endogenous (ATAD3 antibody) and overexpressed (HA antibody) ATAD3A in cells from (B), analyzed by western blots. (D) Working model – mtDNA breaks affect normal cristae morphology through ATAD3A and the MICOS complex (red), hinder protein import, and trigger mitochondrial dysfunction. The ISR is activated to counteract mitochondrial defect, increase mtDNA copy number, and reestablish homeostasis. See also Figure S9.

## Discussion

It is well-established that mtDNA has a shorter half-life than nuclear DNA, which may prevent the accumulation of mutations in the mitochondria (Gross et al., 1969; Shokolenko et al., 2009). DNA lesions, such as oxidized bases or unrepaired nicks, can be converted into DSBs that lead to mtDNA degradation (Suter and Richter, 1999). The recovery of mtDNA copy number necessitates proper communication between the mitochondria and the nucleus. In this study, we used a mitochondrial-targeted restriction enzyme to investigate the cellular response to mtDSBs. A subset of cells with mtDNA breaks displayed severe mitochondrial defects, including a loss of membrane potential, impaired mitochondrial protein import, and defective cristae organization. mtDNA damage also triggered an ISR through the phosphorylation of eIF2α by the DELE1-HRI pathway. We propose that the ISR activation is necessary to counteract the defects in mitochondrial protein import and supports the recovery of mtDNA copy number to restore mitochondrial function (Figure 6D).

Our study revealed that only a small percentage (∼15%) of cells with mtDNA breaks displayed severe defects in their mitochondria and elicited an ISR. One explanation could be that a threshold of mtDNA breaks is necessary to cause the observed mitochondrial defects. However, although the GFP-cells had similar mtDNA copy number, which serves as a proxy for mtDNA breaks (Figure S2F), only the GFP+ cells exhibited membrane depolarization, mitophagy, and loss of IMM proteins (Figure S4A-F). Alternatively, given the transient nature of the ISR induced by mtDSBs, only a subset of cells display import defect at a given time point and are captured as GFP+ cells. In support of this scenario, we show that the GFP signal in GFP+ cells was attenuated over time (Figure 4C and S6C).

Based on our data, the deformation of mitochondrial membranes and loss of cristae junctions are central to triggering a cellular response to mtDSBs and potentially underlie the activation of ISR. Altering mitochondrial membrane integrity and mitochondrial depolarization could hinder the import of DELE1, resulting in its cytoplasmic accumulation and activation of HRI (Horvath et al., 2015). This is consistent with previous literature indicating that mitochondrial import defect results in the accumulation of DELE1 precursor in the cytoplasm and activates the ISR independent of OMA1 (Fessler et al., 2022). The defect could be augmented by the loss of the MICOS complex (Figure S4F) and potential configurational change of ATAD3A, where disruption of the IMM, including the contact sites between IMM and OMM, could further enhance the cytosolic accumulation of DELE1 (Fessler *et al*., 2022; Horvath *et al*., 2015; Shammas et al., 2022).

It has been established that nucleoids reside between cristae and in proximity to the inner mitochondrial membrane (Gerhold et al., 2015; Stephan et al., 2019). Cristae abnormalities associated with mtDSBs recapitulate observations in cells from patients harboring mtDNA mutations (Arbustini et al., 1998), suggesting that the inner mitochondrial membrane could act as a sensor of damaged genomes. We speculate that ATAD3A is the linchpin that connects mtDNA and the IMM. This is consistent with the scaffolding function of ATAD3A, interacting with the membrane organization machinery (OPA1, YME1L, prohibitin, and MICOS complexes) (Arguello et al., 2021) while also associating with nucleoids (He et al., 2012; Ishihara *et al*., 2022; Peralta et al., 2018; Sen *et al*., 2022). Paradoxically, our mass spectrometry data showed an enhanced association of ATAD3A with nucleoids upon mtDNA breaks (Figure 5E). The affinity of ATAD3A to mtDNA may depend on DNA supercoiling. Thus, its binding to relaxed DNA could trigger a conformational shift that would impair the scaffolding role of the AAA+ ATPase. One can also envision a conformational change in ATAD3A bound to linear DNA could interfere with its oligomerization ability. This step was previously implicated in mitochondrial fragmentation and mtDNA stability (Zhao et al., 2019). To that end, future biophysical and structural studies are necessary to elucidate the underpinnings of ATAD3A misfolding in response to defective nucleoids.

Ultimately, while our study uncovered ISR activation in response to exogenously induced mtDNA breaks, our findings are likely to have implications for a broader range of pathological mtDNA perturbations. It has been reported that mutations in Twinkle helicase, which lead to mtDNA replication defect and mtDNA deletions, lead to increased fibroblast growth factor 21 (FGF21), a downstream target of ATF4 (Forsstrom et al., 2019; Khan et al., 2017). Disease models associated with defective exonuclease activity of POLG accumulate linear mtDNA and also show elevated FGF21 (Trifunovic et al., 2004; Wall et al., 2015). In summary, our study adds mtDNA breaks to the repertoire of defects eliciting an ISR in mammalian cells. A better understanding of ISR in mitochondrial DNA diseases could provide potential therapeutic insights.

### Limitation of our study

In our study, we employed mito-ApaLI to introduce mtDNA breaks in an inducible manner and decipher the downstream cellular response. Given the high efficiency of cutting, we were unable to control the exact levels of mtDNA breaks per cell. As a result, we could not determine the exact threshold of mtDNA breaks that would trigger mitochondrial dysfunction. That said, give the the mitochondrial compartmentalization imposed by regulated cristae structures, it is conceivable that a single defective nucleoids could trigger local mitochondrial dysfunction (Jakubke et al., 2021). Nevertheless, better tools to control break formation spatiotemporally are necessary to provide better resolution to the mitochondrial response to mtDNA breaks.

## Acknowledgments

We thank Carlos Moraes for providing the mito-ApaLI plasmid. We acknowledge the Proteomics Laboratory, the Cytometry and Cell Sorting Laboratory, and the DART Microscopy Lab at NYU Langone Health, supported partly by NYU Langone Health and the Laura and Isaac Perlmutter Cancer Center support grant P30CA016087 from the National Cancer Institute. We acknowledge the Integrated Genomics Operation Core at MSKCC, funded by the NCI Cancer Center Support Grant (CCSG, P30 CA08748), Cycle for Survival, and the Marie-Josée and Henry R. Kravis Center for Molecular Oncology. We also acknowledge the Flow Cytometry Core Facility at MSKCC, funded partly through the NIH/NCI Cancer Center Support Grant P30 CA008748. We thank members of the Sfeir lab for their feedback on the manuscript. This work was supported by grants from the Pew Charitable Trust, The G. Harold & Leila Y. Mathers Charitable Foundation, and the Edward Mallinckrodt Jr. Foundation Award (A.S.).

## Authors’ Contribution

A.S. and Y.F. conceived the experimental design and wrote the manuscript. Y.F. performed all experiments. O.S. performed co-immunoprecipitation in Figure S9A. E.K and B.U. performed mass spectrometry and analysis. All authors discussed the results and commented on the manuscript.

## Authors’ information

Agnel Sfeir is a co-founder, consultant, and shareholder of Repare Therapeutics. Correspondence and requests for materials should be addressed to A.S.

## Supplemental Figures

**Figure S1 Related to Figure 1.**
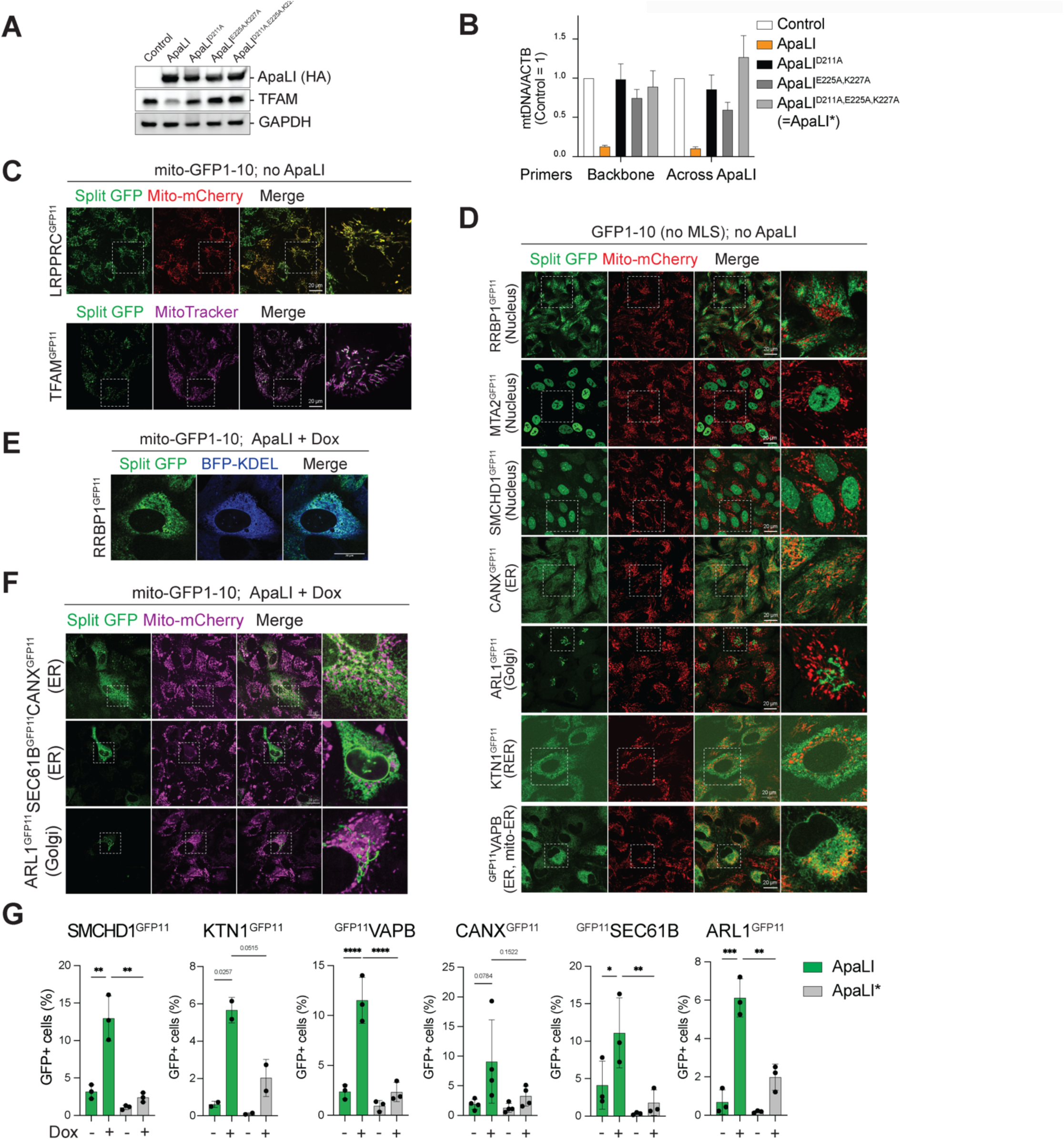
Establish a split-GFP report to assess mitochondrial protein import in response to mtDNA breaks. (A) \Expression of mito-ApaLI (HA antibody) and TFAM in 293T cells transfected with wildtype and catalytically inactive mito-ApaLI for two days, analyzed by western blots. Control sample transfected with a GFP plasmid. (B) mtDNA copy number in 293T cells with the indicated treatments as in (A), measured by qPCR using primers against the backbone (MT-ND2) or across the ApaLI site (MT-RNR1/2) relative to ACTB. All samples were normalized to control cells transfected with a GFP plasmid. (C) Representative image of reconstituted split-GFP in cells expressing mito-GFP1-10 and endogenously GFP11-tagged mitochondrial proteins, LRPPRC (LRPPRC^GFP11^) and TFAM (TFAM^GFP11^). The mitochondrial network is marked by mito-mCherry or MitoTracker staining. (D) Representative images of reconstituted split-GFP in cells expressing the indicated proteins endogenously tagged with GFP11 and a GFP1-10 without MLS. Expression of non-mitochondrial GFP1-10 was used to confirm the proper GFP11 targeting of the genes. The expected localization of the protein being tagged is indicated in parenthesis. RER, rough ER; mito-ER, mitochondria-ER contact site. Scale bar, 20 μm. (E) Representative image of reconstituted split-GFP co-localized with the ER network, labeled with mTagBFP-KDEL, in RRBP1^GFP11^ cells expressing mito-GFP1-10 and mito-ApaLI, and treated with Dox for two days. Scale bar, 20 μm. (F) Representative image of reconstituted split-GFP in cells expressing mito-GFP1-10 and GFP11-tagged CANX (CANX^GFP11^), SEC61B (SEC61B^GFP11^), or ARL1 (ARL1^GFP11^) under mtDSBs. GFP-tagged cells expressing mito-ApaLI were treated with Dox for two days. (G) Percentage of GFP+ population from cells expressing mito-GFP1-10 and indicated GFP11-tagged proteins under mtDSBs, analyzed by flow cytometry. GFP11-tagged cells expressing mito-ApaLI and mito-ApaLI* were treated with Dox for two days. Data are means ± s.d. of *n* = 3 to 4 biological replicates; one-way ANOVA. \**P* ≤ 0.05, \*\**P* ≤ 0.01, \*\*\**P* ≤ 0.001, \*\*\*\**P* ≤ 0.0001.

**Figure S2 Related to Figure 1.**
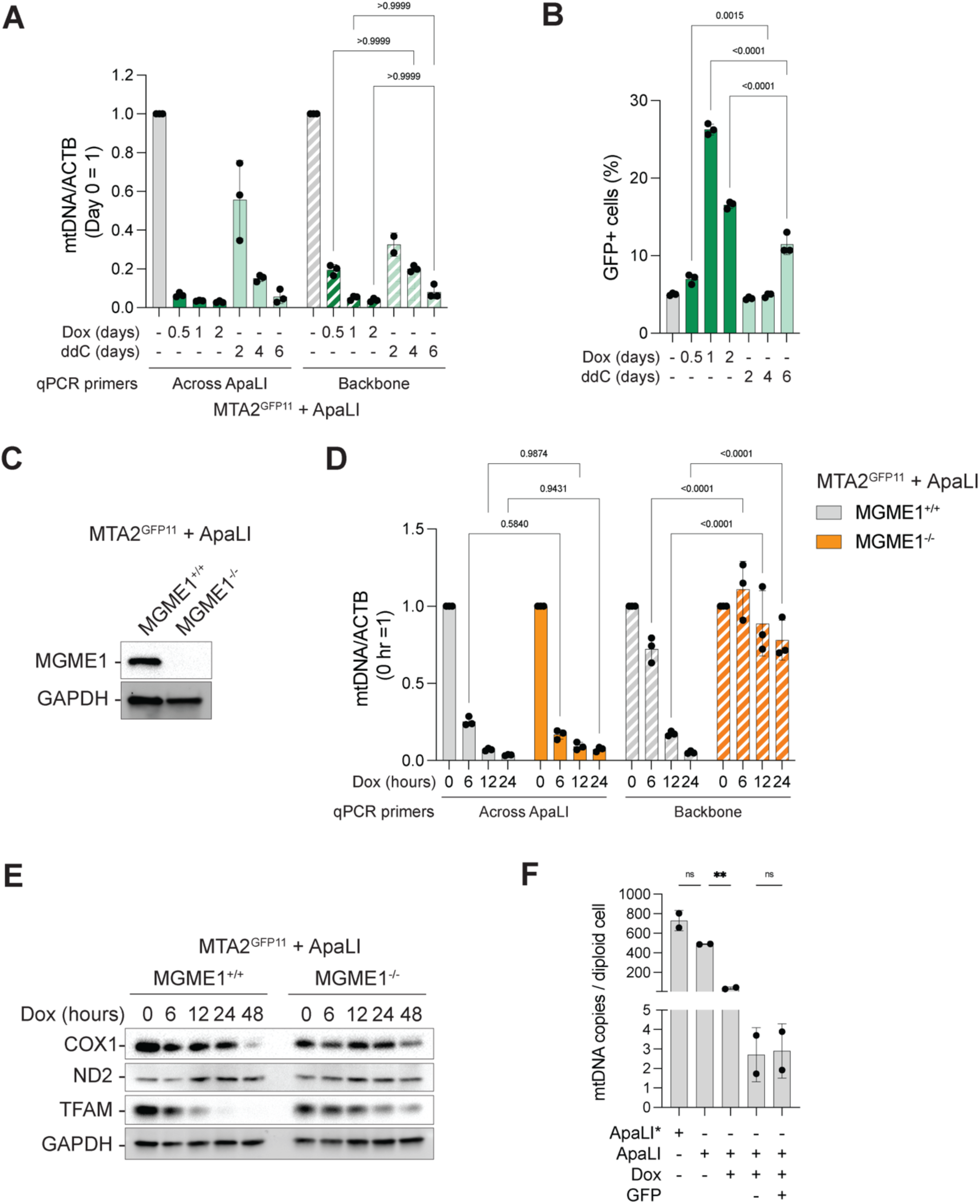
The effect of mtDSBs versus mtDNA depletion in defective mitochondrial protein import. (A) mtDNA copy number of MTA2^GFP11^ cells with inducible mito-ApaLI treated with Dox for 0 – 2 days, or ddC for 0 – 6 days, analyzed by qPCR using primers across ApaLI site (MT-RNR1/2) or against the backbone (MT-ATP6/8) relative to ACTB. Data are means ± s.d. of *n* = 3 biological replicates; two-way ANOVA. (B) Percentage of GFP+ cells from samples in (A), analyzed by flow cytometry. Data are means ± s.d. of *n* = 3 biological replicates; two-way ANOVA. Samples with equivalent remaining mtDNA levels, as shown in (A), were compared. (C) Expression of MGME1 in MTA2^GFP11^ cells where MGME1 was depleted using CRISPR/Cas9 and analyzed by western blots. (D) mtDNA copy number of MGME1-depleted MTA2^GFP11^ cells expressing Dox-induced mito-ApaLI for 0 – 24 hours, analyzed by qPCR using primers across ApaLI site (MT-RNR1/2) or against the backbone (MT-ATP6/8) relative to ACTB. Data are means ± s.d. of *n* = 3 biological replicates; two-way ANOVA. (E) Expression of mtDNA encoded proteins (COX1 and ND2) and TFAM in MGME1-depleted MTA2^GFP11^ cells expressing Dox-induced mito-ApaLI for 0 – 48 hours, analyzed by western blots. (F) mtDNA copy number per cell in sorted GFP-versus GFP+ MTA2^GFP11^ cells under mtDSBs, quantified by digital droplet PCR. Data are means ± s.d. of *n* = 2 biological replicates. One-way ANOVA. **P ≤ 0.01.

**Figure S3 Related to Figure 1.**
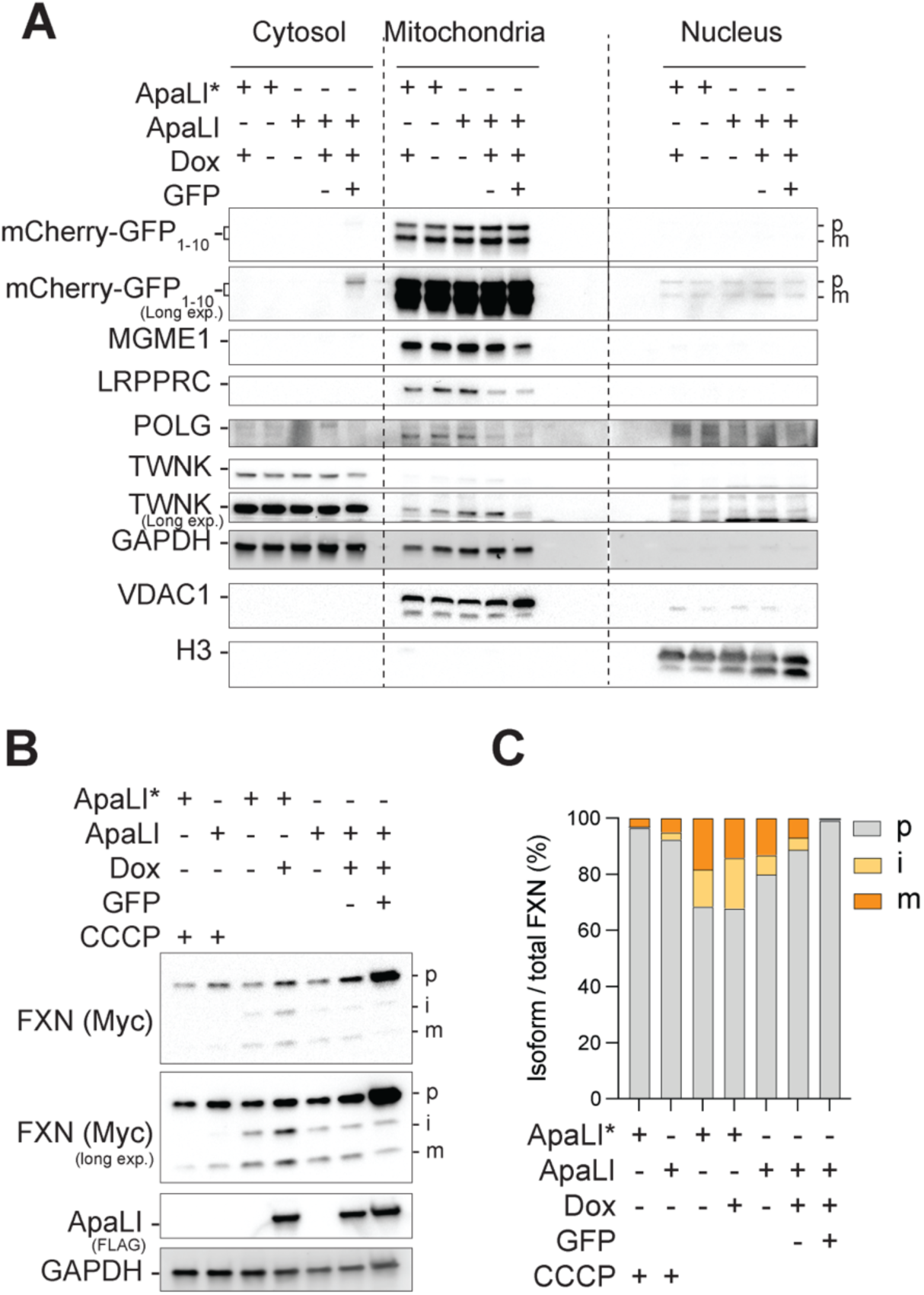
Defective mitochondrial protein import in response to mtDNA breaks. (A) Subcellular fractionation followed by western blot analysis of the indicated proteins from MTA2^GFP11^ cells with indicated treatments, including cell sorting based on GFP. p, precursor isoform; m, mature isoform. Long exp., long exposure. (B) Expression of three isoforms of frataxin (FXN) transfected to sorted GFP- and GFP+ populations of MTA2^GFP11^ cells expressing Dox-induced mito-ApaLI for three days, analyzed by western blots. Mitochondrial uncoupler CCCP was used at 20 μM for 6 hours. i, intermediate isoform. (C) Western blot quantification of frataxin isoforms in (B) normalized to the total level of all FXN isoforms.

**Figure S4 Related to Figure 2.**
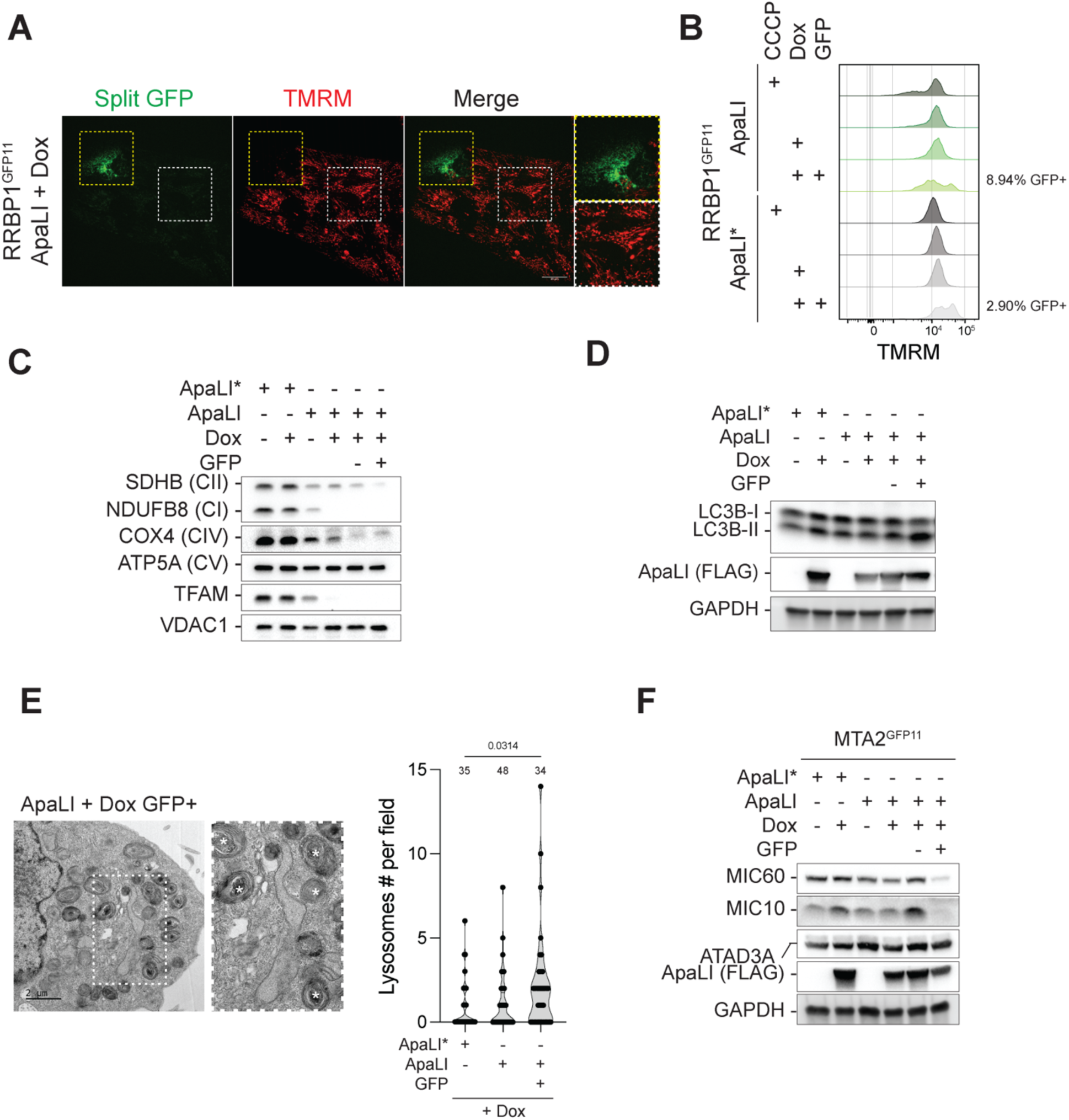
Defective mitochondrial membranes in cells subjected to mtDNA damage. (A) Representative image of TMRM staining in RRBP1^GFP11^ cells expressing mito-GFP1-10 and mito-ApaLI and treated with Dox for two days. Yellow and white boxes depict examples of GFP+ and GFP-cells, respectively. (B) Histograms showing TMRM intensity distribution on the X-axis and unit area on the Y-axis for samples in Figure 2B; the percentages of GFP+ cells are labeled. Mitochondrial uncoupler CCCP was used at 20 μM for 6 hours. (C) Expression of nucleus-encoded OXPHOS proteins in the mitochondrial fraction of GFP- and GFP+ populations from MTA2^GFP11^ cells expressing Dox-induced mito-ApaLI for three days, analyzed by western blots. (D) Expression of LC3B-I and LC3B-II in sorted GFP+ population of MTA2^GFP11^ cells expressing Dox-induced mito-ApaLI for three days, analyzed by western blots. (E) Lysosome accumulation in GFP+ population from cells expressing induced mito-ApaLI. Left, examples of lysosomes marked by (*) in the representative TEM. Right, violin plot presenting lysosome numbers per field. The number of fields analyzed for each sample is labeled above the violins. Two-way ANOVA. (F) Expression of MIC60, MIC10, and ATAD3A in MTA2^GFP11^ cells expressing Dox-induced mito-ApaLI or mito-ApaLI* for three days.

**Figure S5 Related to Figure 3.**
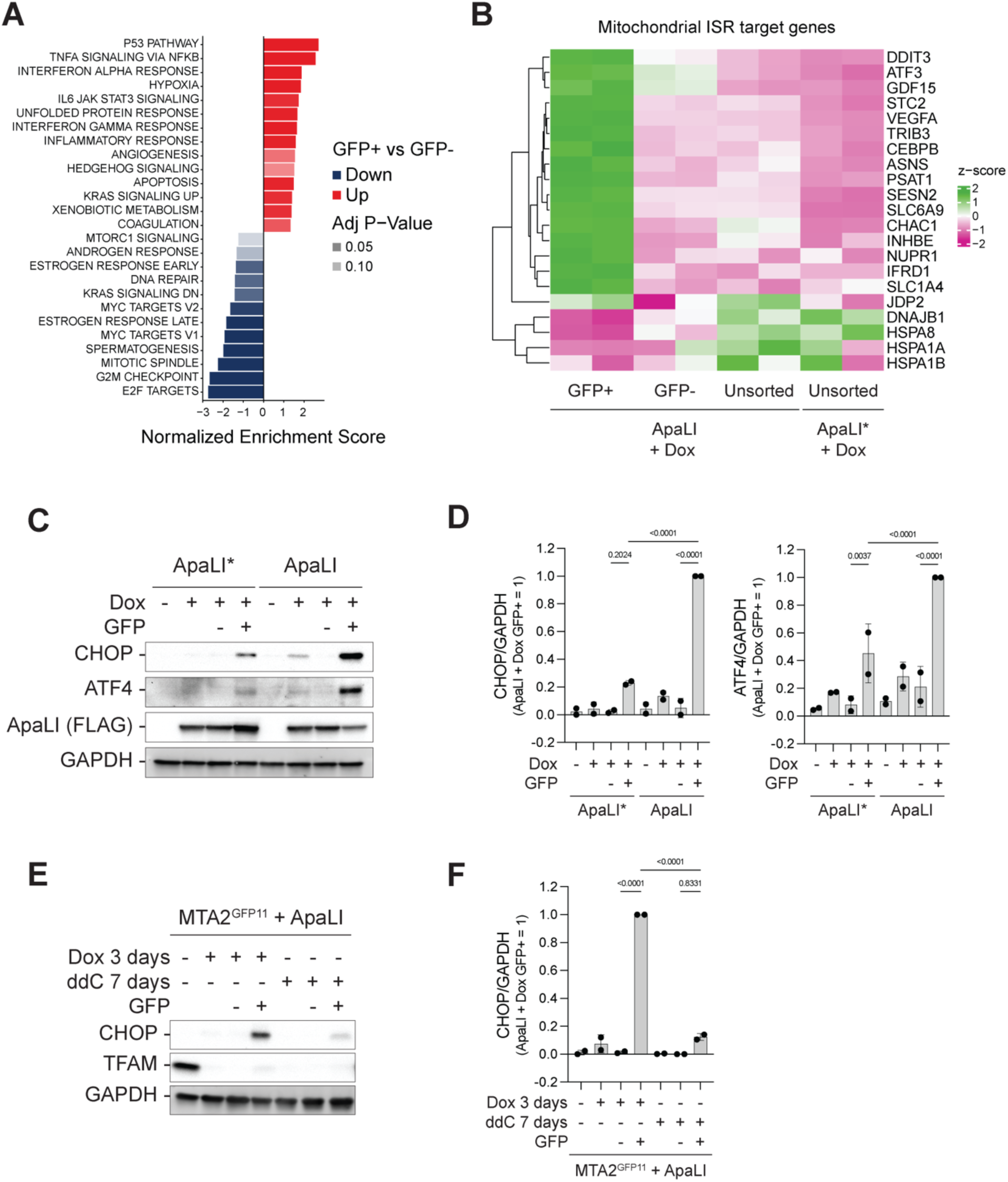
Activation of the integrated stress response in a subset of cells under mtDSBs. (A) Gene Set Enrichment Analysis of differentially expressed genes in GFP+ versus GFP-MTA2^GFP11^ cells induced with Dox for mito-ApaLI expression using MSigDB hallmark gene sets. Red bars indicate up-regulated pathways, and blue bars represent down-regulated genes in GFP+ cells. (B) Heatmap showing the expression of mitochondrial ISR target genes in GFP+, GFP-, and unsorted MTA2^GFP11^ cells expressing mito-ApaLI or mito-ApaLI*. The list of ISR target genes specific to mitochondrial dysfunction was published previously (Fessler et al., 2020). (C) Expression of CHOP and ATF4 in sorted GFP+ populations from MTA2^GFP11^ cells expressing mito-ApaLI versus mito-ApaLI*, analyzed by western blots. Dox induced expression of mito-ApaLI and mito-ApaLI* for 3 days. (D) Western blot quantification of CHOP and ATF4 normalized to loading control GAPDH in (C). *n* = 2. Data are means ± s.d. of *n* = 2 biological replicates. One-way ANOVA. (E) Expression of CHOP in the sorted GFP+ population from MTA2^GFP11^ cells expressing inducible mito-ApaLI and treated with Dox for three days versus treated with ddC for seven days, analyzed by western blots. (F) Western blot quantification of CHOP normalized to loading control GAPDH for samples from (E). *n* = 2. Data are means ± s.d. of *n* = 2 biological replicates. One-way ANOVA.

**Figure S6 Related Figure 4.**
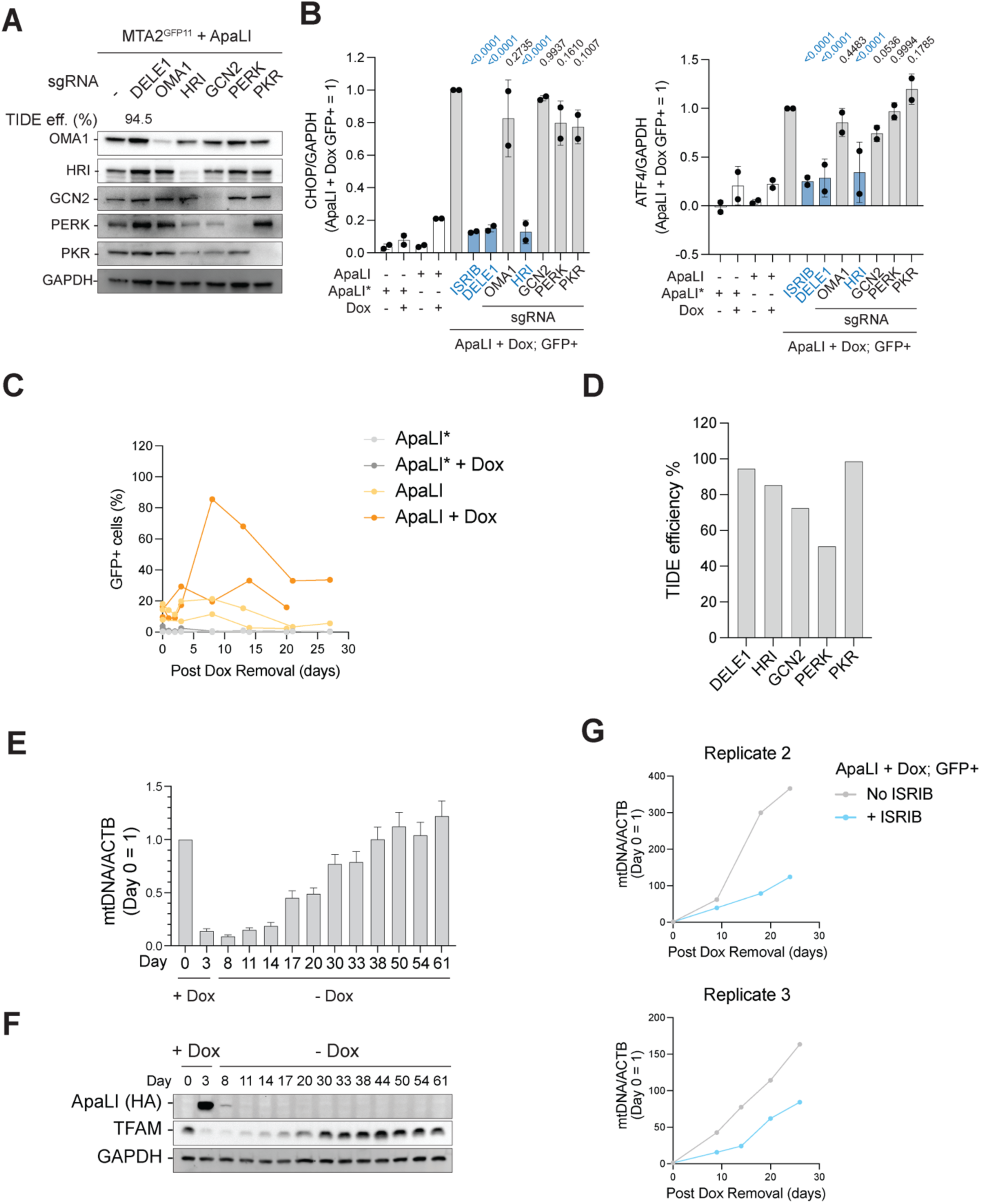
DELE1-HRI activates a robust ISR upon mtDSBs to restore homeostasis. (A) CRISPR/Cas9 editing efficiency in MTA2^GFP11^ cells expressing mito-ApaLI used in Figure 4A. DELE1 was determined by TIDE analysis, and OMA1, HRI, GCN2, PERK, and PRK were assessed by western blots. (B) Western blot quantification of CHOP and ATF4 normalized to GAPDH for samples from Figure 4A and a replicate experiment. Data are means ± s.d. of *n* = 2 biological replicates. Indicated P-values are compared to untreated ApaLI + Dox, GFP+ cells. One-way ANOVA. (C) Temporal dynamics of GFP+ percentage in MTA2^GFP11^ cells expressing Dox-induced mito-ApaLI or mito-ApaLI* for two days, analyzed by flow cytometry. Dox was removed for Dox-treated samples on Day 0. Each line is a replicate. (D) CRISPR/Cas9 editing efficiency analyzed by TIDE in samples represented in Figure 4D. (E) Temporal copy number dynamics of mtDNA upon mtDSBs in ARPE-19 cells, measured by qPCR of mtDNA (MT-ND2) relative to ACTB locus. Cells expressing inducible mito-ApaLI were treated with Dox on Day 0, and Dox was withdrawn on Day 3. Values at different time points were normalized to Day 0. Normalized copy numbers are means ± s.d. from *n* = 3 technical replicates. (F) Expression of ApaLI (HA antibody) and TFAM during mito-ApaLI induction and recovery as in (E), analyzed by western blots. (G) Copy number dynamics of mtDNA in the sorted GFP+ population of MTA2^GFP11^ cells expressing mito-ApaLI and with and without ISRIB treatment Dox was removed post-cell sorting on Day 0. Copy numbers were measured by qPCR using primers against MT-RNR1/2 (mtDNA) relative to ACTB. Values at different time points were normalized to Day 0. See also Figure 4E.

**Figure S7 Related Figure 5.**
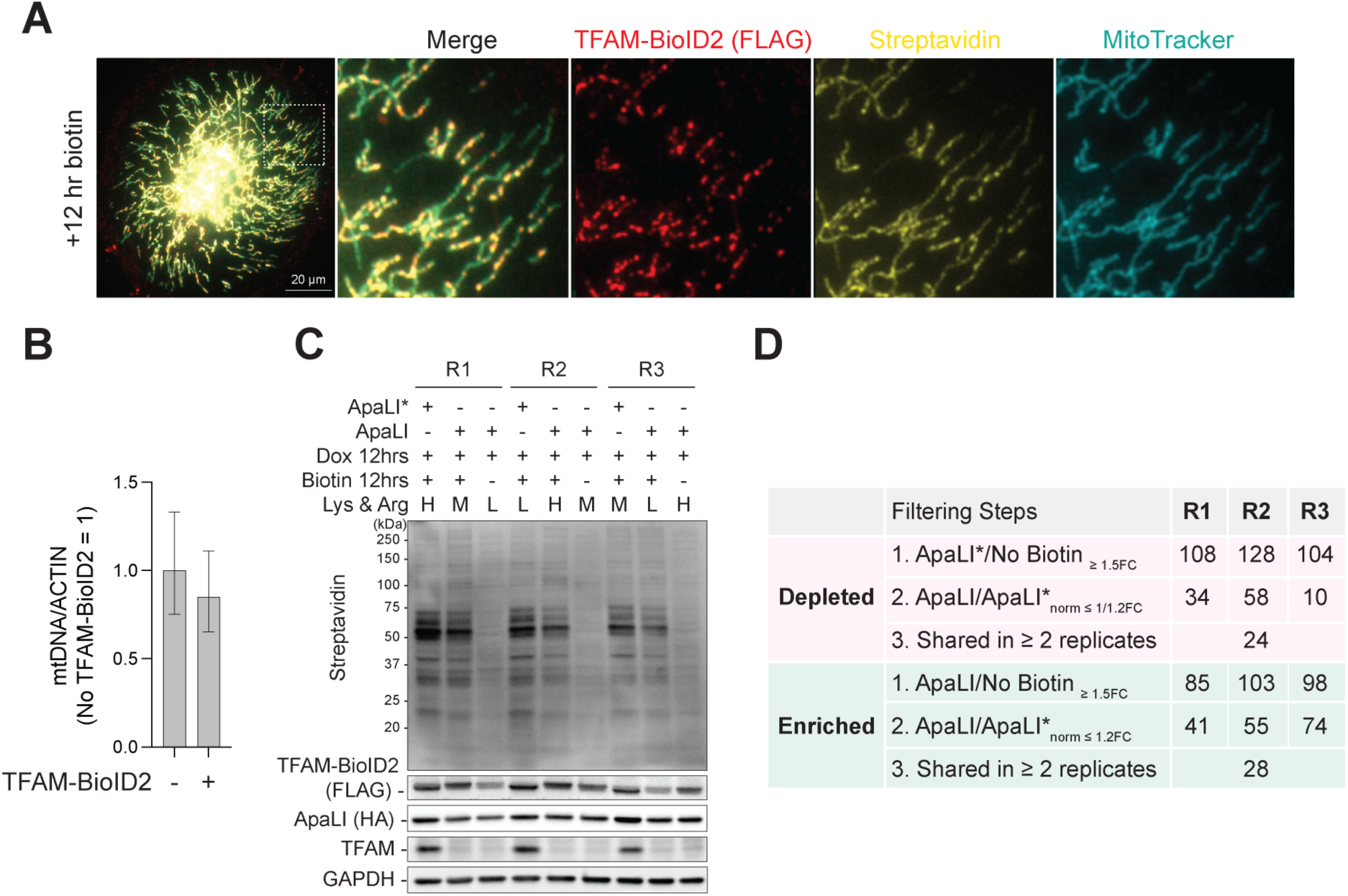
Establishing a TFAM-BioID2-based proteomic approach to identify proteins in the vicinity of nucleoids upon mtDSBs. (A) Representative image of immunofluorescence of TFAM-BioID2 (FLAG antibody) and biotinylated proteins (streptavidin) in ARPE-19 cells expressing TFAM-BioID2 and treated with 50 nM biotin for 12 hours. (B) mtDNA copy number in cells expressing TFAM-BioID2, analyzed by qPCR of MT-RNR1/2 (mtDNA) relative to ACTB. mtDNA levels in cells expressing TFAM-BioID2 were normalized to control cells lacking TFAM-BioID2. Normalized copy numbers are means ± s.d. from *n* = 3 technical replicates. (C) Western blot analysis of protein biotinylation (streptavidin), TFAM-BioID2 (FLAG antibody), and mito-ApaLI (HA antibody) in cell lysates before pooling for streptavidin pull-down and mass spectrometry. Replicates: R1, R2, R3. Lys, lysine; Arg, arginine; H, heavy; M, medium; L, light. (D) Filtering steps to highlight depleted and enriched proteins in cells expressing mito-ApaLI versus mito-ApaLI*. FC, fold change; norm, SILAC ratio normalized to TFAM-BioID2.

**Figure S8 Related to Figure 4.**
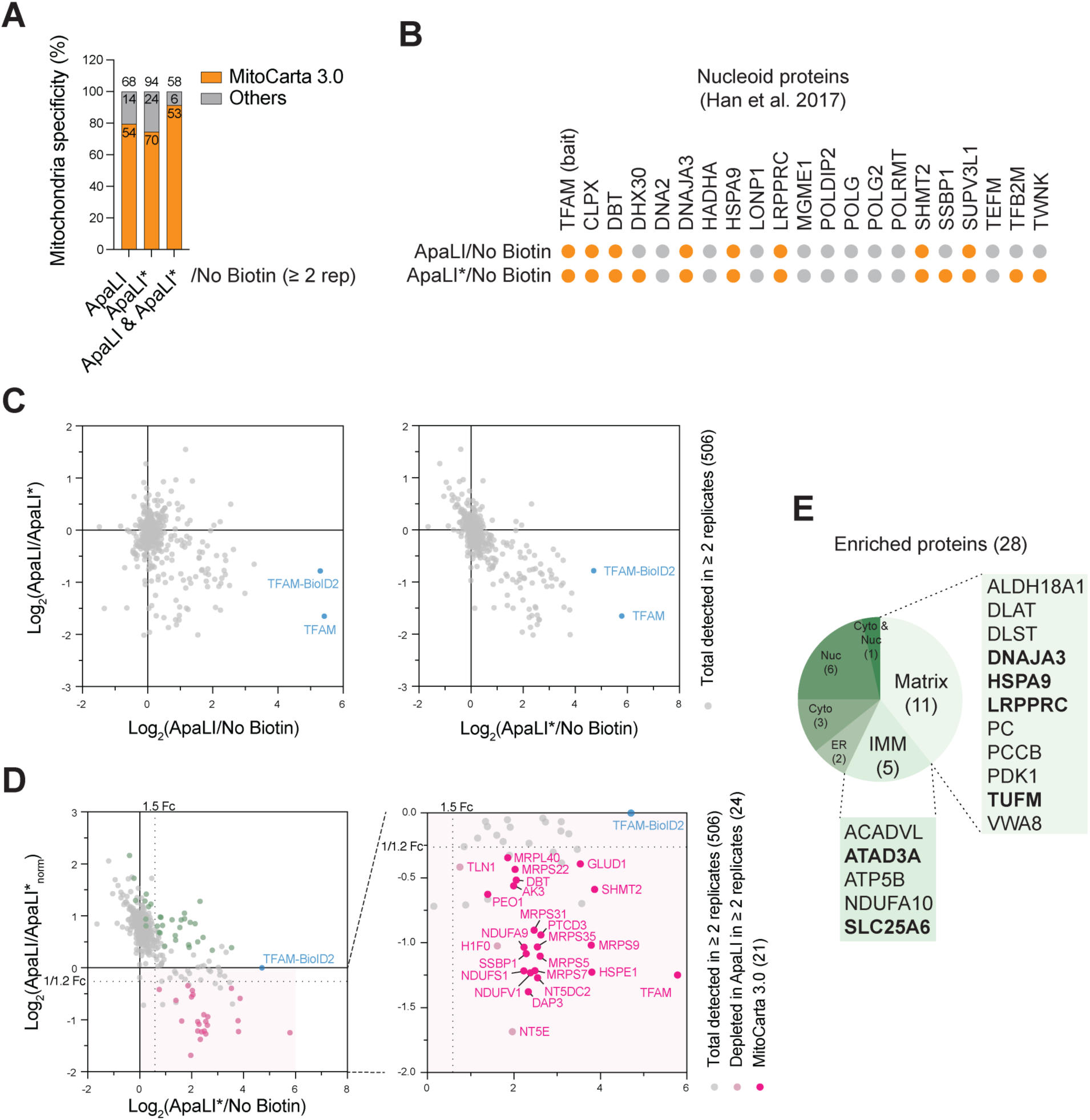
BioID-based proteomic analysis reveals changes in nucleoid composition in response to mtDSBs. (A) Percentage of mitochondrial proteins defined by MitoCarta 3.0 among all proteins captured by TFAM-BioID2. TFAM-BioID2 captured proteins pass Filtering Step 1 in Figure S7D and are present in ≥ 2 replicates. Numbers of proteins are indicated on the bars. ApaLI & ApaLI*, proteins shared between ApaLI and ApaLI*. (B) Nucleoid proteins detected among TFAM-BioID2 captured proteins are labeled in orange. A list of 21 nucleoid proteins was curated (Han et al., 2017). (C) SILAC ratio plot showing the log2(ApaLI/ApaLI*) against log2(ApaLI/No Biotin) or log2(ApaLI*/No Biotin). The SILAC ratios of TFAM-BioID2 were calculated based on peptides from BioID2. (D) SILAC ratio plot showing normalized log2(ApaLI/ApaLI*) against log2(ApaLI*/No Biotin). Twenty-four proteins depleted in mito-ApaLI are highlighted in pink. Among them, 21 MitoCarta3.0 proteins are marked in dark pink and labeled with gene symbols on the right. (E) Categories of subcellular localization of 28 mtDSB-enriched proteins with names listed for those from the mitochondrial matrix and inner membrane (IMM). Proteins in bold were tested in Figure 6A.

**Figure S9 Related to Figure 6.**
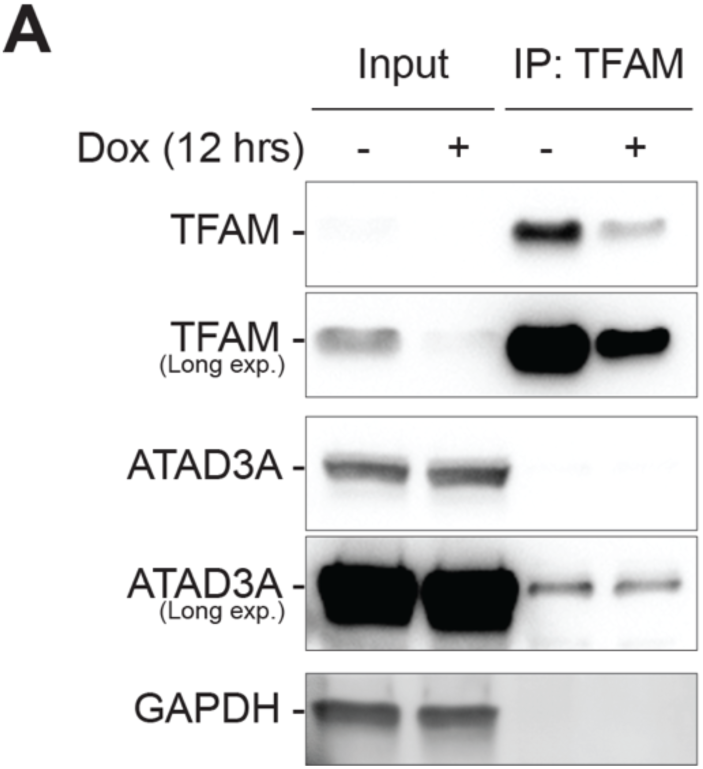
Interaction between ATAD3A and TFAM upon mtDSBs. (A) Co-immunoprecipitation of endogenous TFAM and western blot analysis of TFAM, ATAD3, and GAPDH (negative control) from ARPE-19 cells subjected to Dox-induced ApaLI mtDSBs for 12 hours.

**Table 1** SILAC mass spectrometry results of TFAM-BioID2 associated proteins in response to mtDSBs.

**Table 2** Oligos used in this paper, including primers for qPCR and PCR, CRISPR/Cas9 editing sgRNAs and donors, etc.

**Table 3** Antibodies and dilutions used in western Blots of this paper.

**Video 1** Electron tomogram of mitochondria in MTA2^GFP11^ cells with mito-ApaLI* induced for two days.

**Video 2** Electron tomogram of mitochondria in MTA2^GFP11^ cells with mito-ApaLI induced for two days.

**Video 3** Electron tomogram of mitochondria in GFP+ MTA2^GFP11^ cells with mito-ApaLI induced for two days.

**Video 4** 3D reconstructed mitochondria in MTA2^GFP11^ cells with mito-ApaLI* induced for two days.

**Video 5** 3D reconstructed mitochondria in MTA2^GFP11^ cells with mito-ApaLI induced for two days.

## STAR Methods

### Plasmids

mito-ApaLI (pcDNA3-ApaLI-HA) (Bayona-Bafaluy *et al*., 2005) was generously provided by Carlos Moraes. The catalytically inactive mutants of mito-ApaLI, mito-ApaLI^D211A^, mito-ApaLI^E225A, K227A^, and mito-ApaLI^D211A, E225A, and^ ^K227A^ were generated through overlapping PCR using primers harboring the desired mutations. Doxycycline-inducible mito-ApaLI and mito-ApaLI* plasmids used in Figure 5 and S7-8 (lenti-TRE3G-ApaLI(*)-Hygro) were cloned into LT3REVIR (Addgene Plasmid #111176) by inserting mito-ApaLI (*) (HA-tagged) after a Tet-On 3G promoter and adding Hygromycin selection marker. The doxycycline-inducible mito-ApaLI (*) plasmid (TLCV2-ApaLI(*)-T2A-mTagBFP2-Puro) used in all other figures was cloned using the all-in-one Dox-inducible lentiviral backbone of TLCV2 (Addgene Plasmid #97360) by inserting mito-ApaLI (*) (FLAG-tagged) after the Tet-responsive promoter (TRE), with an addition of a T2A-mTagBFP fragment in-frame with mito-ApaLI (*).

The TFAM-BioID2 fusion plasmid (lenti-CMV-TFAM-BioID2-IRES-Puro) was cloned by first inserting TFAM cDNA into the backbone of MCS-BioID2-HA(Kim *et al*., 2016) (Addgene Plasmid #74224) with the addition of a short linker (2xGGGGS) between TFAM and BioID2. The region containing the CMV promoter and TFAM-BioID2 was cloned into a lentiviral vector, including the Puromycin selection marker.

The lentiviral mito-mCherry-GFP1-10 plasmid (pHAGE2-EF1a-F0ATPMLS-mCherry-GFP1-10) was created by cloning the mitochondria-localized mCherry-GFP1-10 region from plasmid MTS-mCherry-GFP1-10-Hyg-N1(Ruan et al., 2017) (Addgene Plasmid #91957) into a lentiviral backbone with an EF-1α promoter driving the expression. For TMRM staining (Figure 2A, 2B, S4A, and S4B), mCherry was removed from the mito-mCherry-GFP1-10 plasmid by cloning. GFP1-10 plasmid (pHAGE2-EF1a-GFP1-10-IRES-Puro) was generated by cloning GFP1-10 from pcDNA3.1-GFP (1-10) (Addgene Plasmid #70219) into a lentiviral vector that contains EF-1α promoter and Puromycin selection marker. The lentiviral BFP-KDEL plasmid (pHAGE2-EF1a-BFP-KDEL-IRES-Hygro) was based on BFP-KDEL (Addgene Plasmid #49150). Frataxin plasmid (pCMV6-FXN-Myc-FLAG) was purchased from Origene (RC204880). The lentiviral 4xGFP11-SEC61B plasmid (pHAGEF2-EF1a-4xGFP11-SEC61B-IRES-Hygro) was generated by cloning SEC61B from mCherry-SEC61B (Addgene Plasmid #121160), adding 4xGFP11 fragment in-frame to its 5’ end, and inserting the entire 4xGFP11-SEC61B fragment into a lentiviral vector that contains EF-1α promoter and Hygromycin selection marker. The lentiviral HA-ATAD3A plasmid (pHAGE2-EF1a-HA-ATAD3A-IRES-Hygro) was generated by cloning ATAD3A from ATAD3A ORF clone (Genescript #OHU20158D), adding HA tag on N-terminus and inserting into a lentiviral vector containing EF-1α promoter and Hygromycin selection marker.

### Cell culture procedures and treatment

All cells were grown in an incubator at 5% CO_2_ and 37°C. ARPE-19 cells (ATCC CRL-2302) were immortalized with hTERT. ARPE-19 cells and 293T cells (ATCC CRL-3216) used for lentiviral production are cultured in Dulbecco’s modified Eagle medium (DMEM, Corning, 10-013-CV) supplemented with 10% Bovine Calf Serum (GeminiBio), 2 mM L-glutamine (Gibco), 100 U/ml Penicillin-Streptomycin (Gibco), 0.1 mM MEM non-essential amino acids (Gibco), and 50 ug/ml uridine (Sigma-Aldrich). Before the introduction of dox-inducible mito-ApaLI (or mito-ApaLI*) by transfection or lentiviral transduction, cells were switched to be cultured in Dulbecco’s modified Eagle medium (DMEM, Corning, 10-013-CV) supplemented with 10% tetracycline-free fetal bovine serum (FBS, TaKaRa Cat. #631368), 2 mM L-glutamine (Gibco), 100 U/ml Penicillin-Streptomycin (Gibco), 0.1 mM MEM non-essential amino acids (Gibco), and 50 μg/ml uridine (Sigma-Aldrich).

All lentiviral productions were done by transfecting 293T cells with an equal amount of lentiviral packaging combo (pMDLg/pRRE (Addgene Plasmid #12251), pRSV-Rev (Addgene Plasmid #12253), VSV-G (pMD2.G, Addgene Plasmid #12259) in 1:1:1 molar ratio), and plasmid of interest, and changing to fresh media 12 hours post-transfection. Supernatant of 293T cells containing fresh lentivirus was collected every 24 hours after media change and applied to cells to be infected.

Mito-ApaLI and mito-ApaLI* were induced with 1 μg/ml doxycycline (Dox). For mtDNA depletion, 20 μM ddC was added to cells with indicated time and renewed at least every 2 days. For ISR inhibition, ISRIB (Sigma-Aldrich #SML0843) was added to cells at 200 nM for at least 12 hours before mito-ApaLI induction.

Transfection of siRNAs was performed with Lipofectamine RNAiMAX Transfection Reagent (Invitrogen) and following the manufacturer’s protocol.

### Western blot analysis

Cells harvested were counted and lysed in 10 μl Laemmli buffer (4% (w/v) SDS, 20% glycerol, 120 mM Tris-HCl, pH6.8) per 100,000 cells. Lysates were vortexed and boiled at 95°C for 10 minutes. Proteins from equivalent numbers of cells were loaded on SDS-PAGE gels. For mtDNA-encoded proteins (COX1 and ND2) (Figure S2E), cells harvested were lysed in RIPA buffer (150 mM sodium chloride, 1% NP-40, 0.5% sodium deoxycholate, 0.1% SDS, 50 mM Tris, pH 8.0) and sonicated at 4°C. Protein concentration was measured using the Pierce BCA protein assay kit (Thermo Fisher) and equal micrograms of proteins were mixed with Laemmli buffer before being loaded to SDS-PAGE gels. SDS-PAGE gels were transferred to nitrocellulose or PVDF membranes using Trans-Blot Turbo Transfer System or Mini Trans-blot cells (Bio-Rad). Membranes were blocked in 5% milk in TBST (137 mM NaCl, 2.7 mM KCl, 19 mM Tris Base, 0.1% Tween-20) or 5% BSA in TBST for p-eIF2α for 1 hour, followed by incubation with primary antibodies in 1% milk (1% BSA for p-eIF2α) overnight at 4°C. Subsequently, membranes were washed with TBST 3 times, incubated with HRP-linked secondary antibodies at 1:2,500 dilution, and washed again. Membranes were then developed with Clarity Western ECL Substrate (Bio-Rad) and imaged on ChemiDoc XRS+ Imager (Bio-Rad). Loading control a Rhodamine detected GAPDH conjugated antibody against GAPDH. Western blots were quantified using Image lab software (Bio-Rad). A complete list of antibodies and their dilutions used in the study is available in Table 3.

### Genomic DNA isolation

Purified total genomic DNA was used for mtDNA copy number measurement by qPCR and southern blot. Cell pellets harvested were resuspended in 400 μl PBS buffer containing 0.2% (w/v) SDS, 5 mM EDTA, and 0.2 mg/ml Proteinase K and incubated at 50°C for 6 hours with constant shaking at 1,000 rpm. DNA was precipitated by adding 0.3 M Sodium Acetate, pH5.2, and 600 μl isopropanol and kept at -20°C for over 2 hours. Precipitated DNA was centrifuged at 20,000 x *g* at four °C for 30 minutes, followed by a 70% ethanol wash. DNA was then resuspended in TE buffer (10 mM Tris-HCl, pH8.0, 0.1 mM EDTA) and quantified by Nanodrop.

### mtDNA copy number

To determine the relative mtDNA copy number by qPCR, sequences specific to mtDNA (ND2, MT-RNR1/2) and the nuclear ACTB locus (Table 2) were independently amplified from 12.5 ng of total genomic DNA with ssoAdvanced SYBR Green Supermix (Bio-Rad) in a total volume of 10 μl with standard cycling condition suggested by Bio-Rad. qPCR reactions were run on Roche LightCycler 480 or Applied Biosystems QuantStudio 6 Real-Time PCR System. The LightCycler 480 software used the primary relative quantification method with mtDNA as the target and ACTB as a reference. In the QuantStudio 6 software, a Comparative C_T_ experiment was used, with ACTB as the endogenous control.

To determine the absolute mtDNA copy number by digital droplet PCR, we followed the ddMDM method (O’Hara et al. Genome Research. 2019) for cell lysis. Following Proteinase K treatment, samples were diluted with ddH_2_O 1:125 to obtain the final concentration of 10 cell equivalents per μl. 1 µL cell lysate was combined with locus-specific primers, FAM-labeled MT-ND4 (mtDNA) and HEX-labeled EIF2C (nuclear DNA) probes (Table 2), Hind III, and digital PCR Supermix for probes (no dUTP). Cycling conditions for the probes were tested to ensure optimal annealing/extension temperature and optimal separation of positive from empty droplets. Optimization was done with a known positive control. All reactions were performed on a QX200 ddPCR system (Bio-Rad catalog # 1864001), and each sample was evaluated in technical duplicates or triplicates. Reactions were partitioned into a median of ∼10,000 droplets per well using the QX200 droplet generator. Emulsified PCRs were run on a 96-well thermal cycler using cycling conditions identified during the optimization step (95°C 10 seconds; 40 cycles of 94°C 30 seconds and 54, 56, or 60°C 1 second; 98°C 10 seconds; 4°C hold). Plates were read and analyzed with the QuantaSoft software to assess the number of positive droplets for the gene of interest, reference gene, or neither. mtDNA copy numbers per cell were calculated as follows: MT-ND4 droplet number / EIF2C droplet number * 2.

### Southern blot

Purified genomic DNA was digested with BamHI-HF restriction enzyme (NEB) at 5 units per 1 μg DNA for 6 hours at 37°C. Products and Quick-Load 1kb Extend DNA Ladder (NEB) were separated on a 0.6% agarose gel supplemented with ethidium bromide at 30 V for 24 hours at 4°C. The ladders were imaged on ChemiDoc XRS+ Imager (Bio-Rad) before blotting. The agarose gels went through the following washes: 1) depurination (0.25 M HCl) for 30 minutes, 2) twice denaturation (1.5 M NaCl, 0.5 M NaOH) for 30 minutes, 3) twice neutralization (3M NaCl, 0.5 M Tris-HCl, pH7.0) for 30 minutes. The gels were blotted to Amersham Hybond-N membranes (GE) via overnight capillary transfer with 20x SSC (3M NaCl, 0.3 M sodium citrate). The membranes were crosslinked under UV at 1200x100 μJ/cm^2^ and then incubated in pre-hybridization buffer (1.5x SSC, 1% SDS, 7.5% dextran sulfate, and 125 μg/ml salmon sperm DNA) at 65°C for 1 hour. Probes specific to human mtDNA were PCR amplified from 3 regions as in (Moretton *et al*., 2017), and probes specific for nuclear 18S ribosomal DNA were amplified using primers as in (Peeva et al., 2018). For probe labeling, 75 ng of each of the purified PCR products was mixed with 1 μl Random Hexamers (50 μM, Invitrogen) in a total volume of 30 μl, which was then boiled at 95°C for 5 minutes and cooled to room temperature. The reaction was supplemented with 1 μl DNA Polymerase I, Large (Klenow) fragment (NEB), 5 μl dCTP [α-32P] (3000Ci/mmol), 1x NEBuffer 2 and 33 μM dATP, dGTP, dTTP, and incubated at RT for 90 minutes. After adding 50 μl TNES buffer (20 mM EDTA, 400 mM NaCl, 0.5% SDS, 50 mM Tris base), reactions were deactivated at 65°C for 10 minutes. Probes were then purified through a Sephadex G-50 fine (GE) column made with TNE buffer (50 mM Tris-HCl, pH7.4, 100 mM NaCl, 0.1 mM EDTA) in a 3 ml syringe. Eluted probes (in 1 ml TNES buffer) were boiled at 95°C for 5 minutes and immediately added to the membrane in the pre-hybridization buffer. After overnight hybridization at 65°C, membranes were washed once with 4x SSC, three times with 2x SSC supplemented with 0.1% SDS, and wrapped and exposed to storage phosphor screens, which were scanned using Typhoon Imager (GE).

### Immunofluorescence

ARPE-19 cells expressing TFAM-BioID2 were plated on coverslips in 6-well plates and treated with 50 nM biotin for 12 hours. Cells were stained with 125 nM MitoTracker Deep Red (Invitrogen) for 30 minutes. Cells were rinsed with PBS buffer and fixed with 4% paraformaldehyde for 10 minutes at RT. After PBS washes, cells were permeabilized with 0.5% Triton X-100 in PBS buffer for 10 minutes at RT, followed by PBS washes. Cells on coverslips were blocked with blocking buffer (1 mg/ml BSA, 3% goat serum, 0.1% Triton X-100, 1mM EDTA) for 30 minutes at RT and incubated with FLAG antibody (Sigma F3165, 1:10000) and Alex Fluor 488 conjugated Streptavidin (Invitrogen S11223, 1:2000) in blocking buffer overnight at 4°C. Cells were then washed with PBS and hybridized with Alexa Fluor 594 conjugated anti-mouse secondary antibody for 1 hour at RT. After PBS washes and DAPI staining, coverslips were mounted on glass slides with ProLong Gold Antifade mountant (Invitrogen). Slides were imaged on a Nikon ECLIPSE Ei2-E inverted microscope.

### Tagging of endogenous proteins and gene knockout by CRISPR/Cas9

For tagging of genes of interest with GFP11, the CRISPR ribonucleoprotein (RNP) complex of gRNA and Cas9 protein (IDT), together with a single-stranded donor (ssODN), were delivered into mito-GFP1-10 cells using the Nucleofector system (Lonza). Cell line constitutively expressing mito-mCherry-GFP1-10 was generated by transducing ARPE-19 cells with lentiviral mito-mCherry-GFP1-10 and enriched for mCherry positive cells via cell sorting. gRNAs were designed close to the stop codon of the genes (and the start codon of VAPB). 500 nt ssODNs containing MYC and 4xGFP11 flanked by ∼100 nt homology arms and mutated PAMs (protospacer adjacent motif) were ordered from IDT. A complete list of gRNA targets and ssODNs used in the study is available in Table 2. RNP complexes preparation was based on IDT protocol by 1) annealing an equal amount of 100 μM Alt-R CRISPR-Cas9 crRNA and 100 μM tracrRNA to form gRNA and 2) incubating 150 pmol gRNA with 125 pmol Alt-R Cas9 enzyme for 20 minutes. 5 μl RNP complex,

1.5 μg ssODN, and 1.2 μl 100 μM Alt-R Cas9 Electroporation Enhancer (IDT) were mixed and added to 20,000 cells resuspended in 18 μl SF Nucleofection Buffer (Lonza). The Nucleofection mixture was transferred to 16-well Nucleocuvette strips (Lonza) and nucleofected with the 4D-Nucleofector X unit (Lonza) using the ER-100 program. After recovering in 100 μl RPMI 1640 media (Gibco) for 10 minutes, cells were plated in prewarmed culture media supplemented with 1 μM Alt-R HDR Enhancer V2 (IDT). After 48-72 hours, cells were plated at clonal density in 15 cm plates for clonal cell isolation. Colonies were screened by genotyping PCR using primers flanking the editing site (outside of homology arm regions) and looking for larger PCR products potentially containing GFP11 insertion. Clonal cells with potential homozygous or heterozygous knock-in were chosen and further confirmed by transducing GFP1-10 without MLS and detecting GFP signal at the primary localization of the protein (e.g. ER for RRBP1, VAPB, KTN1; golgi for ARL1). Confirmed clones were introduced with mito-ApaLI and mito-ApaLI* by lentiviral transduction to examine the GFP signal upon mtDSBs. For GFP11 tagging of SEC61B, the ARPE-19 cell line expressing mito-mCherry-GFP1-10 was transduced with lentiviral 4xGFP11-SEC61B and selected for integration using Hygromycin. Confirmation of proper localization of SEC61B and introduction of mito-ApaLI and mito-ApaLI* was done the same way as above for cells knocked in with GFP11.

Gene knock-outs followed the same procedure of CRISPR RNP delivery, excluding ssODN and HDR enhancer. For Figure S2C, MTA2^GFP11^ cells with inducible mito-ApaLI were nucleofected with CRISPR RNA against the MGME1 gene. For Figure 4A and 4D, MTA2^GFP11^ cells were nucleofected with CRISPR RNP. Knock-out efficiencies mainly were confirmed by western blots, except that DELE1, which has no available antibodies (Guo *et al*., 2020), was confirmed by PCR (Table 2) and TIDE analysis (https://tide.nki.nl). MTA2^GFP11^ cells with ISR factor depletion were expanded and transduced with mito-ApaLI. Cells were induced with Dox and sorted for GFP+ cells for downstream western blot analysis. For Figure 6A, 6B and S9A, MTA2^GFP11^ cells with inducible mito-ApaLI were nucleofected with CRISPR RNP. 48-72 hours after CRISPR delivery, cells were induced with Dox for 2 days and analyzed by flow cytometry and western blots.

### Flow cytometry

For flow cytometry analysis of split GFP signal, GFP11 tagged cells expressing Dox-induced mito-ApaLI or mito-ApaLI* (-T2A-mTagBFP) were trypsinized, centrifuged, resuspended in 3% FBS in PBS, and passed through cell strainers. Cells were analyzed on an LSR Fortessa analyzer (BD) with GFP signal detected through 488 nm laser and 525/50 Bandpass (BP) filter, mCherry detected through 561 nm laser and 610/20 BP filter, mTagBFP detected through 405 nm laser and 450/50 BP filter. Recorded data were analyzed using FlowJo software. After gating for singlet living cells, mTagBFP negative populations were gated for samples without Dox induction, and mTagBFP positive cells were gated for samples with Dox-induced. Subsequently, GFP+ populations were gated based on mito-ApaLI* without Dox treatment, and the same gating was applied across all samples.

### Live cell imaging by spinning disk confocal microscope

All imaging on split GFP signal was performed at live cell imaging conditions. Cells were plated on glass bottom plates (Cellvis) and induced with Dox. Before imaging, media was changed to live cell imaging media (DMEM, high glucose, HEPES, no phenol red (Gibco, 21063029) supplemented with 10% Bovine Calf Serum (GeminiBio), 2 mM L-glutamine (Gibco), 100 U/ml Penicillin-Streptomycin (Gibco), 0.1 mM MEM non-essential amino acids (Gibco), and 50 ug/ml uridine (Sigma-Aldrich)). Cells were imaged on a Nikon CSU-W1 Spinning Disk Confocal microscope with a 100x oil lens. TOKAI HIT STX system maintained cells at 37°C, 5% CO_2,_ and humidity. Within every experiment, laser percentage and exposure time for each channel were set the same across all samples in comparison. Captured images were then analyzed on Fiji software by setting the same minimum and maximum intensity for samples in comparison.

### Subcellular fractionation

Cells were harvested and processed using Cell Fractionation Kit (Abcam ab109719) following standard protocol with optimization for ARPE-19 cells. 50 μl 1x Buffer A and 50 μl Buffer B (or Buffer C) were used for 500,000 cells. Buffer B was prepared by diluting Detergent I 500-fold in 1x Buffer A. Buffer C was prepared by diluting Detergent II 12.5-fold in 1x Buffer A. Cytosol, and mitochondrial extractions were both done on a rotator for 30 minutes at RT. All centrifugation steps during the extraction were done for 5 minutes at 4°C.

### Frataxin transfection

MTA2^GFP11^ cells expressing mito-ApaLI and mito-ApaLI* were induced with Dox for 48 hours. MTA2^GFP11^ cells with mito-ApaLI induction were sorted for GFP- and GFP+ populations by flow cytometry. 350,00 cells per sample were seeded in a 6-well to recover overnight. Cells were transfected with 0.5 μg frataxin plasmid using Lipofectamine 3000 (Invitrogen) and 36 hours later harvested for western blot analysis. Treatment of 20 μM CCCP was done 6 hours before harvesting.

### TMRM staining and analysis

ARPE-19 cells were CRISPR/Cas9 targeted to introduce 4xGFP11 to the C-terminal of RRBP. The confirmed homozygous clone was transduced with mito-GFP1-10 (without mCherry), followed by transduction of mito-ApaLI and mito-ApaLI*. Cells were plated in glass-bottom plates (Cellvis). After being induced with Dox for 2 days, cells were stained with 1 μM TMRM (Invitrogen) in culture media for 30 minutes at 37°C. Subsequently, media was replaced with live cell imaging media as indicated in Live cell imaging. For flow cytometry analysis of the TMRM signal, cells were trypsinized, centrifuged, and resuspended in 3% FBS in PBS after staining. Cells were analyzed on an LSR Fortessa analyzer. TMRM signal was detected through a 561 nm laser and 586/15 Bandpass filter. Recorded data were analyzed using FlowJo software. TMRM^low^ gating was set based on TMRM-stained mito-ApaLI* cells and was applied to all conditions, including GFP+ and GFP-subpopulations. mito-ApaLI and mito-ApaLI* cells were treated with 20 μM CCCP for 6 hours as a positive control. While mito-ApaLI* cells without Dox induction were resistant to CCCP, mito-ApaLI* cells were sensitive to membrane potential loss upon CCCP treatment.

### RNA sequencing (RNA-seq) and analysis

Four conditions were prepared for the RRBP1^GFP11^ (or MTA2^GFP11^) cell line: 1) mito-ApaLI* treated with Dox (unsorted), 2) unsorted, 3) GFP-subset, 4) GFP+ subset of mito-ApaLI treated with Dox, with two replicates for each condition. For each sample, 500,000 cells were harvested as a pellet and kept frozen at -80°C before handing over to the Integrated Genomics Operation at MSKCC for RNA purification, TruSeq stranded mRNA (polyA) library preparation, and sequencing.

Frozen cells were lysed in 1 ml TRIzol Reagent (Thermo Fisher 15596018), and phase separation was induced with 200 µl chloroform. According to the manufacturer’s protocol, RNA was extracted from 350 µl of the aqueous phase using the miRNeasy Mini Kit (Qiagen 217004) (on the QIAcube Connect (Qiagen). Samples were eluted in 33 µl RNase-free water. After RiboGreen quantification and quality control by Agilent BioAnalyzer, 500 ng of total RNA with RIN values of 8.9 - 10 underwent polyA selection and TruSeq library preparation according to instructions provided by Illumina (TruSeq Stranded mRNA LT Kit, RS-122-2102), with 8 cycles of PCR. Samples were barcoded and ran on a NovaSeq 6000 in a PE100 run, using the NovaSeq 6000 S4 Reagent Kit (200 Cycles) (Illumina). An average of 45 million paired reads was generated per sample. The percent of mRNA bases averaged 91%.

RNA-seq data were analyzed using an automated pipeline Seq-N-Slide (10.5281/zenodo.5550459.) designed to work on the BigPurple HPC cluster at NYU Langone Health. RNA-seq analysis of the Seq-N-Slide pipeline used STAR (Spliced Transcripts Alignment to a Reference) for alignment, DESeq2 for differential gene expression analysis, and GSEA (Gene Set Enrichment Analysis) for pathway analysis.

Heatmaps in Figure 3A, 3D, and S5B present the z-scores of individual genes, which were calculated by applying regularized log (`rlog()`) and scaling (`scale()`, across all 8 samples) functions to the `.rds` file derived from DESeq2 analysis. The list of mitochondrial ISR target genes used in Figure 3D and S5B was obtained from Fessler, et al. Nature. 2020 (Fessler *et al*., 2020) and is composed of ATF4, CHOP target genes, and heat shock protein genes that are induced by CCCP and are dependent on DELE1–HRI–eIF2α signaling.

### Transmission electron microscopy and analysis

Cultured cells were fixed in 0.1 M sodium cacodylate buffer (pH 7.4) containing 2.5% glutaraldehyde and 2% paraformaldehyde for 2 hours and post-fixed with 1% osmium tetroxide and 1% potassium ferrocyanide for one hour at 4°C, then block stained in 0.25% aqueous uranyl acetate, processed in a standard manner, and embedded in EMbed 812 (Electron Microscopy Sciences, Hatfield, PA). 70 nm ultrathin sections were cut (UC6 microtome; Leica Microsystems) and mounted on 200 mesh copper grids. 200 nm thickness sections were collected on slotted copper grids (Electron Microscopy Sciences, Hatfield, PA) coated with a formvar membrane. All sections were counterstained by incubation with 3% uranyl acetate in 50% methanol for 20 min, followed by washing in water and incubation with Renyld’s lead citrate for 5 min.

For Figure 2C and S4E, 70 nm ultrathin sections were imaged under Talos120C transmission electron microscope (Thermo Fisher Scientific) with Gatan (4k x 4k) OneView Camera (Gatan, Inc., Pleasanton, CA). 30-50 fields per sample were randomly imaged at 8000x magnification for statistical analysis. For cristae morphology quantification, mitochondria sections with paralleled and lamellar cristae were considered “normal,” and mitochondria sections without paralleled and lamellar cristae, including complete loss of cristae, were considered as “missing/aberrant” cristae morphology.

200 nm thickness sections were used for the tomogram in Figure 2E and Video 1-5. In detail, 10 nm gold particles were put on grids as fiducial markers before a thin layer of carbon (Auto306 Vacuum Evaporator; Edwards BOC) was evaporated on top of 200 nm sections to minimize beam-induced specimen shrinkage during data collection for tomograms. Samples were imaged with JEOL 1400 Flash TEM (JEOL Ltd.) equipped with Gatan Rio16 camera at 8,000X nominal magnification (corresponding to a 1.27-nm pixel size) and 2.1 μm defocus. Dual-axis tilt series were collected with a dual-axis tomography holder (Model 2040, Fischione Instruments) using a tilt range of ± 65° with a 2° increment (SerialEM 3.0). To collect the tilt series about the second axis, the sample was rotated 90° using the holder’s sample rotation mechanism, and the same region was reimaged. Fiducial alignment of projection images and calculation of 3-D volumes was carried out using Protomo software (Winkler and Taylor,2006), and dual-axis reconstruction was created in IMOD (Kremer et al., 1996; Mastronarde, 1997). Reconstructions were manually segmented in ORS Dragonfly (The Objects) after applying a 3 × 3 × 3 median filter.

### Proximity-based protein biotinylation and streptavidin pull-down

ARPE-19 cells were transduced with lenti-TRE3G-ApaLI(*)-Hygro lentivirus and selected with 300 μg/ml Hygromycin. Due to the high heterogeneity of mito-ApaLI(*) expression among cells after Dox induction, we picked clonal cells that were confirmed with 100% mito-ApaLI(*) induction efficiency by immunofluorescence (IF). Two clones with a equivalent expression of mito-ApaLI and mito-ApaLI* upon Dox treatment were chosen for TFAM-BioID2 introduction. mito-ApaLI and mito-ApaLI* cells were transduced with a relatively low level of TFAM-BioID2 lentivirus and selected with 1 μg/ml Puromycin. IF confirmed proper TFAM-BioID2 localization and biotinylation after treating the cells with 50 nM biotin.

To metabolically label the proteomes of TFAM-BioID expressing mito-ApaLI(*) cells with heavy (H), medium (M), or light (L) isotopes of lysine and arginine, we cultured each cell line in three 6 cm plates starting from 10% confluency. One plate was cultured in light SILAC media: DMEM deficient in L-lysine and L-arginine (DMEM for SILAC, Thermo Fisher, A33822) supplemented with 84 mg/ml L-arginine (Arg0) and 146 mg/ml L-lysine (Lys0) (Sigma-Aldrich), 10% dialyzed fetal bovine serum (Gibco), 100 U/ml Penicillin-Streptomycin (Gibco), 0.1 mM MEM non-essential amino acids (Gibco), and 50 ug/ml Uridine (Sigma-Aldrich). The second plate was cultured in medium SILAC media with the same composition as above, except Arg0 and Lys0 were replaced by L-Arginine [^13^C_6_] HCl (Arg-6) and L-lysine-4,4,5,5-*d*_4_ (Lys-4) (Cambridge Isotope Laboratories). The third plate was cultured in heavy SILAC media with the same composition, except arginine and lysine were replaced by L-Arginine [^13^C_6_,D_7_,^15^N_4_]HCl (Arg-17) and L-Lysine [^13^C_6_,^15^N_2_]HCl (Lys-8) (Cambridge Isotope Laboratories). Cells were passaged and expanded in the corresponding SILAC media for ten doublings.

Each replicate is comprised of three conditions: No Biotin (ApaLI treated with Dox), mito-ApaLI (ApaLI treated with Dox and biotin), and mito-ApaLI*\** (ApaLI* treated with Dox and biotin) (Figure S7C). Within the exact replicate, each condition utilized cells labeled in different SILAC media. Each condition across different replicates also used cells labeled in different SILAC media (Figure S7C). In short, the design of conditions in three replicates with different SILAC isotopes is as follows: Replicate 1, No Biotin (L), ApaLI (M), ApaLI* (H); Replicate 2, No Biotin (M), ApaLI (H), ApaLI* (M); Replicate 3, No Biotin (M), ApaLI (H), ApaLI* (M). Forty million cells were plated in 4 15-cm plates for each condition in each replicate. For biotinylation and mito-ApaLI activation, 50 nM biotin and 1 μg/ml doxycycline were added to the media simultaneously for 12 hours.

Cell lysis and streptavidin pull-down procedures were done as Step 34-41 in Hung et al. Nature Protocol. 2016 (Hung et al., 2016), with mild modifications as follows. Cell pellets of 40 million cells were lysed in freshly prepared 800 μl of RIPA buffer (50 mM Tris, 150 mM NaCl, 0.1% SDS, 0.5% sodium deoxycholate, 1% Triton X-100, 1x Halt Protease and Phosphatase Inhibitor Cocktail (Thermo Fisher), 1 mM PMSF, 10 mM sodium azide, and 10 mM sodium ascorbate) and rotating at 4°C for 30 minutes. Lysates were clarified by centrifugation at 15,000 x *g* for 10 minutes at 4°C. Protein concentration in the supernatant was measured using the Pierce BCA protein assay kit (Thermo Fisher). Whole-cell lysates (5 mg of protein) from the H, M, and L samples of the exact replicate were mixed in a 1:1:1 H:M:L ratio by protein mass. 15 mg of lysate mixtures were incubated with 1.875 ml streptavidin magnetic beads (Dynabeads MyOne Streptavidin C1, Thermo Fisher, 65001) slurry pre-washed with RIPA buffer twice. The suspensions were set at 4°C overnight with gentle rotation. Streptavidin beads were washed twice with 2 ml RIPA buffer, twice with 2M urea in 10 mM Tris-HCl pH 8.0, twice with 2 ml RIPA buffer, and twice with 50 mM Tris-HCl pH7.4, 50 mM NaCl. Biotinylated proteins were eluted by incubating the beads with 225 μl Laemmli buffer (4% (w/v) SDS, 20% glycerol, 120 mM Tris-HCl, pH6.8) supplemented with 20 mM DTT and 2 mM biotin, and heating to 95°C for 5 minutes.

### Protein digestion and extraction

Proteins were reduced with 2.5 μl 0.2 M dithiothreitol (Sigma) for one hour at 57 ⁰C at pH 7.5. Subsequently, samples were alkylated with 2.5 μl 0.5 M iodoacetamide (Sigma) for 45 minutes at room temperature in the dark. NuPAGE LDS Sample buffer (1X) (Invitrogen) was added to the samples, and the samples were transferred to a NuPAGE 4-12% Bis-Tris Gel 1.0 mm x 10 well (Invitrogen) for SDS PAGE gel electrophoresis. The gel was stained with GelCode Blue Stain Reagent (Thermo Scientific). Sample lanes were excised and destained with 1:1 (v/v) methanol and 100 mM ammonium bicarbonate at 4°C with agitation. Destained gel pieces were dehydrated in a SpeedVac concentrator. Dried gel pieces were resuspended in 300 μL 100 mM ammonium bicarbonate with 250 ng Promega trypsin for overnight digestion. A 300 μL solution of 5% formic acid and 0.2% trifluoroacetic acid (TFA) R2 50 μm Poros (Applied Biosystems) beads slurry in water was added to the gel pieces before returning the samples to the shaker for an additional three hours at 4°C. Beads were loaded onto equilibrated C18 ziptips (Millipore), with 0.1% TFA, using a microcentrifuge for 30 seconds at 6,000 rpm. The beads were washed with 0.5% acetic acid. Peptides were eluted with 40% acetonitrile in 0.5% acetic acid, followed by 80% acetonitrile in 0.5% acetic acid. The organic solvent was removed using a SpeedVac concentrator. The samples were reconstituted in 0.5% acetic acid and stored at -80°C until analysis.

### Liquid chromatography (LC) – mass spectrometry (MS)

LC separation was performed online on EASY-nLC 1000 (Thermo Scientific) utilizing Acclaim PepMap 100 (75 um x 2 cm) precolumn and PepMap RSLC C18 (2 um, 100A x 50 cm) analytical column. Peptides were gradient eluted from the column directly to Orbitrap Q Exactive HF-X mass spectrometer using a 215 min acetonitrile gradient from 5 to 26 % B in 179 min followed by the ramp to 40% B in 20 min and final equilibration in 100% B for 15 min (A=2% ACN 0.5% AcOH / B=80% ACN 0.5% AcOH). The flow rate was set at 200 nl/min. High-resolution full MS spectra were acquired with a resolution of 45,000, an AGC target of 3e6, a maximum ion injection time of 45 ms, and a scan range of 400 to 1500 m/z. Following each full MS scan, 20 data-dependent HCD MS/MS scans were acquired at the resolution of 15,000, AGC target of 1e5, maximum ion time of 120 ms, one Microscan, 2 m/z isolation window, NCE of 27, fixed first mass 150 m/z and dynamic exclusion for 30 seconds. Both MS and MS^2^ spectra were recorded in profile mode.

### Analysis to determine the differential proteins in response to mtDSBs

MS data were analyzed using MaxQuant software version 1.6.3.4 (Cox and Mann, 2008) and searched against the SwissProt subset of the human uniprot database (http://www.uniprot.org/) containing 20,430 entries with sequences for mito-ApaLI and BioID2 appended. Database search was performed in Andromeda (Cox et al., 2011b) integrated into the MaxQuant environment. A list of 248 common laboratory contaminants included in MaxQuant was added to the database, and reversed versions of all sequences were. The enzyme specificity was set for searching to trypsin, with the maximum number of missed cleavages set to 2. The precursor mass tolerance was set to 20 ppm for the first search used for non-linear mass re-calibration (Cox et al., 2011a) and then to 6 ppm for the main search. Methionine oxidation was searched as a variable modification; carbamidomethylation of cysteine was explored as a fixed modification. SILAC labeling with 3 channels (referred to as SILAC ratio) was used as a quantification option. The false discovery rate (FDR) for peptide, protein, and site identification was set to 1%; the minimum peptide length was set to 6.

Only proteins with quantified SILAC ratios across all conditions (ApaLI/No Biotin, ApaLI*/No Biotin, ApaLI/ApaLI*) and in ≥ 2 replicates were retained for analysis. We noticed that TFAM-BioID2 (quantified using peptides assigned to BioID2) was reduced in ApaLI compared to mito-ApaLI*\** due to mtDNA degradation after DSBs, similar to TFAM (including endogenous TFAM, and TFAM from TFAM-BioID2 overexpression) (Figure S8C). Given that decreased level of TFAM-BioID2 bait likely leads to a reduction in protein biotinylation, we normalized the ApaLI/ApaLI* SILAC ratio of all proteins to that of TFAM-BioID2.

Proteins depleted upon mtDSBs were regarded as enriched in ApaLI*\** compared to ApaLI. We first ensured proteins captured by TFAM-BioID2 in ApaLI*\** are above the background, using a cut-off of ApaLI*/No Biotin ≥ 1.5 (Figure S7D, Depleted Step 1). Next, we retrieved proteins depleted upon mtDSBs using criteria normalized ApaLI/ApaLI* ≤ 1/1.2 and shared in ≥ 2 replicates (Figure S7D, Depleted Step 2-3).

For proteins enriched upon mtDSBs, we first identified proteins captured by TFAM-BioID2 in ApaLI as the above background, using a cut-off of ApaLI/No Biotin ≥ 1.5 (Figure S7D, Enriched Step 1). Next, we retrieved proteins enriched upon mtDSBs using criteria normalized ApaLI/ApaLI* ≥ 1.2 and shared in ≥ 2 replicates (Figure S7D, Enriched Step 2-3).

### Co-immunoprecipitation (Co-IP)

ARPE-19 cells expressing inducible mito-ApaLI were treated with or without doxycycline for 12 hours. Cells were harvested via trypsinization, washed once with 1X PBS, pelleted and lysed in 1mL of IP Lysis Buffer (50mM Tris-HCl pH 7.5, 150mM NaCl, 0.5% NP-40, 0.1% sodium deoxycholate, 5% glycerol, 1mM EDTA) supplemented with 1x Halt Protease and Phosphatase Inhibitor Cocktail (Thermo Fisher), and 1mM DTT. Cell lysates were sonicated for 5 minutes and incubated on ice for 1 hour. Cell lysates were cleared by centrifugation at the maximum speed at 4°C for 30 minutes. Protein was quantified using Pierce BCA protein assay kit (Thermo Fisher). Around 4 mg of protein was subjected to IP using either 2 μg TFAM antibody (Santa Cruz sc-376672) overnight while rotating at 4°C. 60 μl of Protein G Sepharose 4 Fast Flow (Cytiva 17061801) pre-blocked in 5% BSA were added to each IP sample and incubated for an additional 6 hours while rotating at 4°C. Beads were washed five times in 1 ml of Wash Buffer (50 mM Tris-HCl at pH 7.4, 150 mM NaCl, 1% Triton X-100, 0.1% SDS). Proteins were eluted from the beads in Laemmli buffer by boiling for 5 minutes at 95°C and subjected to western blots.

